# ADEP1 activated ClpP1P2 macromolecule of *Leptospira*, an ideal Achilles’ heel to deregulate proteostasis and hamper the cell survival

**DOI:** 10.1101/2020.08.05.237438

**Authors:** Anusua Dhara, Md Saddam Hussain, Shankar Prasad Kanaujia, Manish Kumar

## Abstract

The caseinolytic protease (ClpP) complex in *Leptospira interrogans* is unusual in its functional activation. The genus *Leptospira* has two ClpPs, ClpP1 and ClpP2, which transcribes independently, regardless it couples to form the active tetradecamer. Acyldepsipeptide (ADEP) antibiotic hampers the growth of numerous bacterial species by activating the target protein ClpP and dysregulating the physiological proteostasis within the cell. *In vitro* culture of the *L. interrogans* fortified with the ADEP impeded the spirochete growth accompanied by a more elongated morphology. The chemoactivation of the ClpP is conditional on the duration of the self-compartmentalization of each of the ClpP isoforms. The small extent (10 min) self-assembled ClpP1P2 revealed inhibition in the peptidase activity (7-fold) in the presence of the ADEP due to the self-cleavage of the ClpP subunits. On supplementation of the β-casein or bovine serum albumin, the peptidase activity of the ClpP1P2 (short-incubated) got enhanced by the ADEP, while the ClpP1P2 (long-incubated) activity was retained to the same level. ADEP can also switch on the ClpP1P2 from a strict peptidase into proteolytic machinery that discerns and degrades the unfolded protein substrates autonomous of the cognate chaperone ClpX. In consensus to the most prokaryotes with the multi ClpP variants, the computational prototype of the ClpP1P2 tertiary structure infers that the hydrophobic pocket wherein the ADEPs predominantly docks are present in the ClpP2 heptamer. Additionally, the dynamic light scattering and the site-directed mutagenesis of a catalytic serine residue in either of the ClpP isoforms proposes a second interaction site for the ADEP.

## INTRODUCTION

*Leptospira interrogans* is the causative agent of leptospirosis, a globally important zoonotic disease (*1*). The transmission of the pathogenic *Leptospira* between animals, humans, and the environment is essential for the maintenance of its enzootic cycle (*2*). Over a million cases of leptospirosis are reported every year, with approximately 60000 deaths in humans(*3*). Leptospirosis being a zoonotic disease disable livestock production in developing tropical and sub-tropical countries where animal rearing is a primary source of livelihood (*4*). Antibiotics, particularly of the penicillin group, are considered as the first-line therapy for leptospirosis (*5*). However, due to the emergent multi-drug resistance of the Gram-negative and Gram-positive bacteria, an urgent need for therapeutics acting on novel pathways to curtail such persistent bacteria is the need of the hour (*6*). The subcellular pathways which are central to the survival of the bacteria during the infection are attractive candidates for new drug design. In such an effort, the acyldepsipeptides (ADEPs), a new class of antibacterial compound and its derivative were found to target the caseinolytic protease (ClpP protease), the proteolytic core of bacterial ATP-dependent proteases (*7, 8*). ADEP1 is a natural molecule of the acyldepsipeptide family produced by *Streptomyces hawaiiensis* that function by dysregulating/activating the ClpP in other microbes unlike other conventional antibiotics (*7, 9, 10*). Activation of the ClpP results in the inhibition of cell division, imbalance in cellular proteostasis, and finally, the cell death of the bacteria including *Staphylococcus*, *Streptococcus*, *Mycobacterium* (*11, 12*). Also, prokaryote ClpP has been found to have a crucial role in regulating processes such as stress tolerance, virulence, morphological differentiation and antibiotic resistance (*10, 13*–*17*). Dysregulating the activity of the Clp protease in the pathogenic bacteria by the ADEP’s or other activators leads to a reduction of its chance for cell survival. The exploitation of such targets is now helpful to destroy multi-drug resistance or the persister form of bacteria emerging due to the improper use of antibiotics (*10, 14, 18, 19*).

Caseinolytic protease system in prokaryotes is composed of the core ClpP catalytic components, regulatory chaperones (ATPases), and the adaptor protein (*20, 21*). Most bacterial species, including *E. coli*, *Bacillus subtilis*, and *Staphylococcus aureus* have one *clpP* gene that, along with their associated ATPases, are nonessential for cell viability whereas, in actinobacteria and cyanobacteria, two or more copies of *clpP* are found, and at least one functional copy is indispensable for viability (*22, 23*). In *E. coli*, the core catalytic component ClpP is a tetradecameric barrel-shaped serine peptidase with the 14 active sites contained within its proteolytic chamber (*24*). In *Mycobacterium tuberculosis*, *clpP1* and *clpP2* form an operon and both the genes product are critical to compose an operative peptidase by stacking the ClpP1 and the ClpP2 homoheptamers into a heterotetradecamer (*22*). It is demonstrated that in *E. coli*, the core ClpP independently can degrade smaller peptides; however, it needs to associate with its cognate Clp/Hsp100 chaperone (Clp-ATPase) to degrade the larger polypeptides and proteins (*25*). The cognate chaperones coordinate with the ClpP in substrate recognition, unfolding of the substrate using energy derived from the ATP hydrolysis and the delivery of the unfolded polypeptide into a proteolytic compartment of the ClpP (*26*). The chaperone ClpX self-composes into a hexamer and employs its peptide loops (IGF/L) to anchor into the apical site (hydrophobic pocket) of the ClpP tetradecamer and render the opening of the entrance pore to foster access of larger substrates in a coordinated strategy (*27*). It has been ascertained that in bacteria with single ClpP isoform, a total of two ClpX or ClpA hexamers can bind to one ClpP barrel from both sites, resulting in a ClpX-ClpP-ClpX or ClpA-ClpP-ClpA complex formation (*28, 29*). Whereas, in bacteria like the *Mycobacterium*, *Listeria* and *Chlamydia* with the multi-ClpP isoforms, the cognate ATPase chaperone has been documented to dock exclusively to the ClpP2 hydrophobic pocket (*30-34*). Biochemical studies in the *B. subtilis* infer the antibiotic ADEP1 mimics ClpX peptide loops and thereby broadens the entrance pores of the ClpP protease and could degrade larger polypeptides unaided as an unregulated protease in the absence of any unfoldase (*35, 36*). In addition to the widening of the entrance pores of the proteolytic compartment, ADEP stabilizes the ClpP tetradecamer and stimulates the catalysis allosterically (*36, 37*).

The Clp protease of the bacteria in association with the ATPase chaperone/unfoldase is a physiological prerequisite for the quality control of the cytosolic proteins (*38*). Manipulating the Clp protease (ClpP) function has exhibited to impact the virulence and infectivity of several different pathogens as discussed in an elegant review elsewhere (*39*). During the late 90s and early 21^st^ century, the ClpP and its allied chaperones were established to have a direct connection with the virulence or stress in the *Staphylococcus aureus* (*13*, *40*, *41*), *Streptococcus pneumoniae* (*42*–*44*), *Listeria monocytogenes* (*45*–*47*) and *Salmonella typhimurium* (*48*–*50*). In a later term, the operating role of the ClpP was determined in a few other microbes like *Pseudomonas aeruginosa* (*51, 52*), *Legionella pneumophila* (*53*), and *Chlamydia* (*54, 55*).

To date, a significant investigation of the ADEPs has been restricted in the bacterial ClpPs in the phylum of the firmicutes, actinobacteria, or the chlamydiae (*26, 56, 57*). In spirochetes, a phylum that encompasses a catalog of pathogenic bacteria like *Leptospira*, *Borrelia*, and *Treponema*, the influence of the ADEPs on its ClpP is yet to be unveiled. In *Leptospira*, the core ClpP catalytic element exists in two isoforms ClpP1 (LIC11417) and ClpP2 (LIC11951) and is transcribed independently (*21*). In the same analysis, the *Leptospira* ClpP1 was ascertained to self-assemble into a larger molecule (14-21mer) than the pure ClpP2 (14-mer), and both of the pure isoforms were functionally dormant (*21*). Nonetheless, the two ClpP isoforms of the *Leptospira* jointly self-assembles into a heterotetradecamer structure composed of two stacked homoheptamer of the ClpP1 and ClpP2 to constitute operative peptidase machinery (*21*). Therefore in this investigation, we explored the influence of the antibiotic ADEP1 on the live *Leptospira* carrying the operative ClpP target by a bacterial growth inhibition assay. The inhibitory impact of ADEP1 on the *Leptospira* growth is validated by various *in vitro* biochemical reactions of recombinant ClpP proteins on model substrates and the computational modeling.

## RESULTS

### ADEP1 treatment elongated *Leptospira* and hampered its growth kinetics

We examined the effect of antibiotic ADEP1 on the growth and morphology of the pathogenic *Leptospira* for a period of 120 h under *in vitro* condition. The measured length of the bacteria under the microscope appeared slightly elongated within the 24 h of sub-culturing the leptospires in a media supplemented with the 10 μg mL-1 ADEP1 (15 μM) (Figure 1A). The average length of the *Leptospira* cells treated with the ADEP1 ranged from 12.8-14.6 μm, whereas the untreated cells were 10.9-11.7 μm. In the presence of ADEP1 (15 μM), there was a 1.2-fold increase in the length of the leptospires. During the ADEP1 (15 μM) treatment period (24 - 120 h), the measured increase in the length of the bacterium was essentially identical under the given *in vitro* condition. The effective bactericidal concentration of ADEP1 for *Leptospira* was assessed by the growth kinetics measurement in the presence of the increasing concentration of ADEP1 (20 - 60 μM). As compared to untreated cells, the growth kinetics of the ADEP1 treated spirochete curtailed from the 48 h onwards in proportion to the amount of ADEP1 (Figure 1B). The decline phase of the spirochete growth curve was achieved within the 48 h of adding 43 μg mL^−1^ ADEP1 (60 μM). The morphology of the spirochetes grown in the presence of a higher ADEP1 concentration (60 μM) was evaluated under electron microscopy and compared with the untreated ones under *in vitro* condition (Figure 1C). The average length of a segment (per three complete spiral turns) of the spirochete treated with ADEP1 was longer (1.2 fold) than the untreated one, implying a reduction in spiral frequency as the rationale behind the spirochete elongation. To understand the toxic effect of ADEP1 on *Leptospira*, we examined the effect of ADEP1 on its target protein ClpP by conducting various *in vitro* biochemical analysis using the recombinant functional protein and computational modeling.

**Figure 1.**
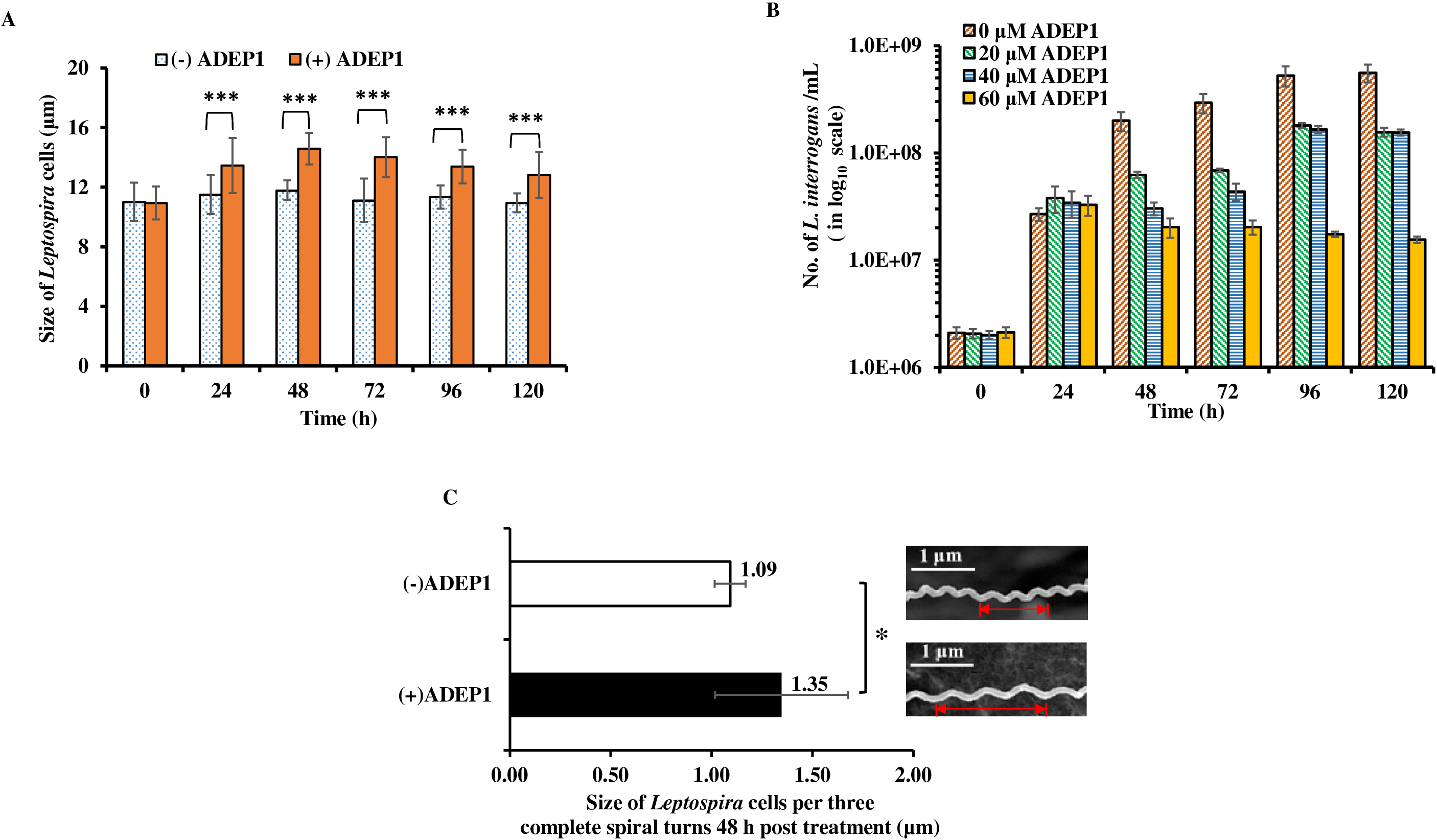
The growth curve and the length of the pathogenic *Leptospira* are affected when grown in the presence of the ADEP1 under *in vitro* condition. **(A)** Bar graph demonstrating the morphology of the *Leptospira* cells was elongated when grown in the presence of the ADEP1 after 24 h of the growth curve. **(B)** Inhibition of the growth curve of the *Leptospira* in the presence of an increasing concentration of the ADEP1 under the *in vitro* condition. The growth of the *Leptospira* was inhibited in the presence of the ADEP1 at 48 h compared to the control. Each experiment was performed independently twice with the three replicates in each set. **(C)** Field emission scanning electron microscopy to measure the morphology change of the *Leptospira* grown in the presence of the ADEP1 under the *in vitro* condition. The average partial length (three complete spiral turns) of the *Leptospira* in the presence (+) of the ADEP1 (60 μM) at the 48 h measured under the FESEM is graphically represented. The representative magnified *Leptospira* image (25 kX resolution and 1 μm scale) treated (+) with the ADEP1 appears to be more relaxed than the untreated one, with an estimated 1.23-fold elongation. The error bars represent the standard errors of the mean (SEM) from the two-independent experiments performed. Student’s t-test was performed for the statistical analysis (***p-value<0.001 and * p-value<0.05).

### ADEP1 bound ClpP1P2 of *Leptospira* triggers autoproteolysis under *in vitro* condition

In our recent study, we demonstrated that the rClpP1P2 peptidase activity of the *Leptospira* is conditional on the ClpP self-assembly duration (*21*). The absolute peptidase activity of rClpP1P2 of *Leptospira* generated after a short-incubation (1 h) for self-assembly was lower than that of long-incubated (24 h) rClpP1P2 (*21*). The explanation for such difference in activity was apparent through the native-PAGE and the dynamic light scattering (DLS) analysis, wherein a more stable and functional population of the rClpP1P2 tetradecamer complex was formed after the long-incubation. In this study, we assessed the peptidase activity of the pure rClpP isoforms or their mixture (rClpP1P2) in the presence and absence of the ADEP1 (5 - 40 μM) towards the model fluorogenic dipeptide substrate S1 (Suc-LY-AMC) used elsewhere (*21, 58*). We generated the rClpP1P2 heterocomplex by mixing the pure rClpP isoforms under the short- (10 min) and the long-incubation period (24 h) before assessing the peptidase activity of the operative heterocomplex in the presence of ADEP1. There was a 7-fold inhibition in the peptidase activity of the rClpP1P2 (short-incubated) in the presence of the ADEP1 (40 μM) versus the basal activity without the ADEP1 (Figure 2A). In contrast, the peptidase activity of the rClpP1P2 (long-incubated) got stimulated by 2.5-fold in the presence of the ADEP1 (15 - 40 μM) than its basal activity without the ADEP1 (Figure 2A). The absolute peptidase activity of the ClpP1P2 (short-incubated) in the presence or absence of the ADEP1 was lower than that of the ClpP1P2 (long-incubated) (Figure S1A, Figure S1B, and Figure S1C). Notably, the relative decline in the peptidase activity of the rClpP1P2 (long-incubated) was observed at the 20 - 40 μM of ADEP1 versus the optimal 15 μM ADEP1 concentration. ADEP1 (up to 40 μM) on the other hand, failed to stimulate any peptidase activity in the pure rClpP isoforms of the *Leptospira* (data not shown). The antibiotic ADEP1 is known to activate the peptidase activity of the ClpP1P2 heterocomplex by an increase in the diameter for the substrate entry into the peptidase machinery. To comprehend the unusual effect of the ADEP1 on the rClpP1P2 (short-incubated) of the *Leptospira*, we resolved the reaction products of the peptidase reaction after completion on a denatured polyacrylamide gel. The staining of the polyacrylamide gel illustrated self-degradation of the rClpP1P2 (short-incubated) subunits in proportion to the ADEP1 supplemented for the peptidase activation (Figure 2B, upper panel). Whereas, the pure rClpP isoforms did not show any degradation in the presence of the ADEP1 (Figure 2B, middle and lower panel). Incredibly, on a polyacrylamide gel, the reaction product of the rClpP1P2 (long-incubated) demonstrated even more self-degradation of the ClpP subunits (Figure 2C). Such self-degradation of the rClpP1P2 (long-incubated) did not reconcile with respect to the gain in the peptidase activity of the rClpP1P2 (long-incubated) in the presence of the ADEP1 (Figure 2A) and motivated us to look forward to other conceivable means of chemoactivation. Hence, ADEP1 mediated functional gain in the rClpP1P2 triggers autoproteolysis of the ClpP protomers, and the chemoactivation of the rClpP1P2 is dependent on the duration self-compartmentalization process of the ClpP isoforms under the given *in vitro* condition.

**Figure 2.**
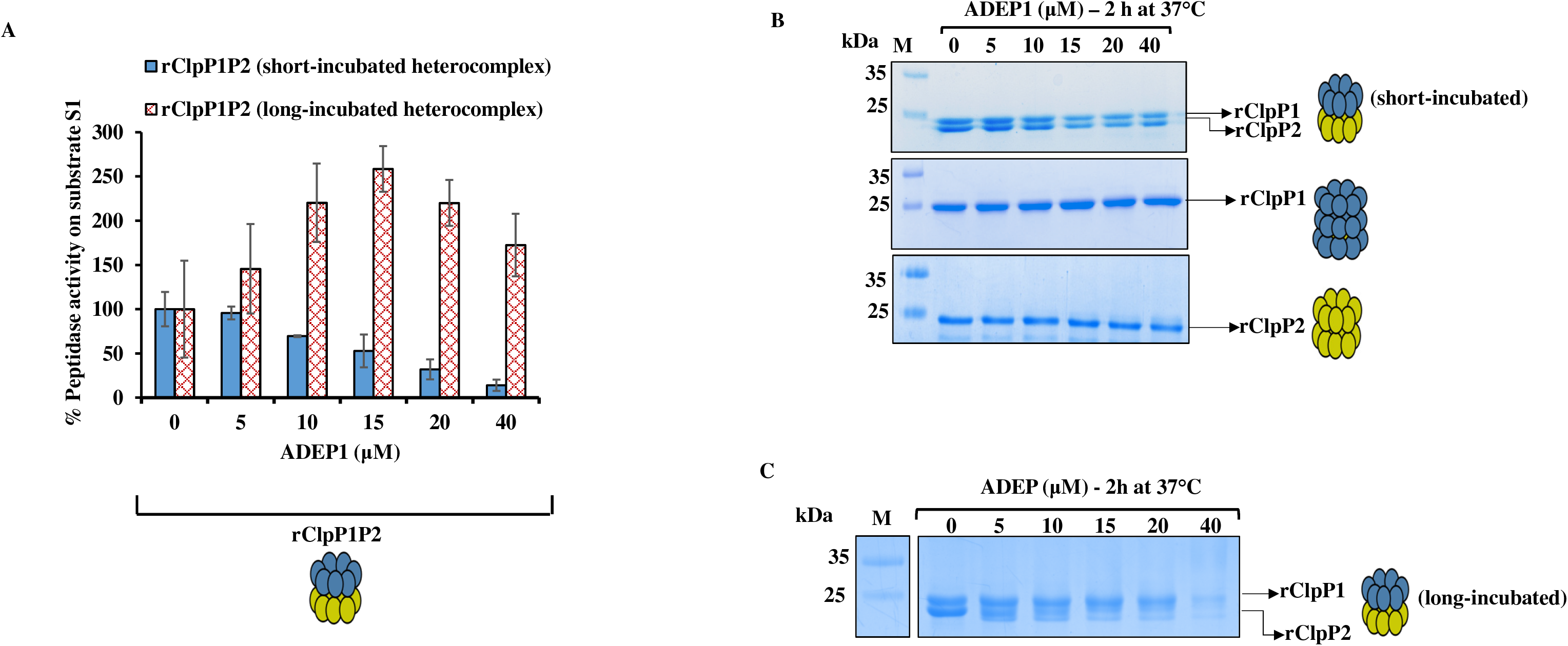
Stimulation of the peptidase activity of the recombinant ClpP1P2 (rClpP1P2) heterocomplex in the presence of the ADEP1 triggers auto proteolysis. **(A)** Peptidase activity of the short-(10 min) and the long-incubated (24 h) rClpP1P2 on the small fluorogenic peptide substrate in the presence of the ADEP1. Peptidase activity of the rClpP1P2 is represented as a percentage (%), wherein the end-point fluorescence was measured after the 2 h of the enzymatic reaction. The measured end-point fluorescence value of the rClpP1P2 (containing no ADEP1) as control was considered as 100% for measuring the relative peptidase activity. The error bars represent the standard errors of the mean (SEM) from the two-independent experiments performed. **(B)** Denaturing polyacrylamide gel electrophoresis of the pure rClpP isoforms and their heterocomplex, rClpP1P2 after ADEP1 treatment. **(C)** Denaturing polyacrylamide gel electrophoresis of the rClpP1P2 (long-incubated) after the ADEP1 treatment. For clarity, the schematics of the ClpP1 and the ClpP2 are represented where each blue and yellow sphere depicts a monomeric subunit of the ClpP1 and the ClpP2, respectively.

### ADEP1 increases the peptidase activity of ClpP1P2 (short-incubated) in the presence of casein or bovine serum albumin and switch the ClpP1P2 to ATPase independent proteolytic machinery

Within bacteria, under the natural conditions, it is unrealistic to develop conditions with a paucity of the protein substrates to the ADEP1 activated ClpP peptidase machinery. Thus, to mimic the natural subcellular ambiance of the bacteria where no dearth of the substrates/proteins for the activated ClpP exists, we modified the peptidase activity assay strategy by introducing an additional β-casein (unstructured substrate) or BSA (bovine serum albumin) protein. On supplementation of the β-casein substrate, the relative peptidase activity of the ADEP1 (15 μM) bound rClpP1P2 (short-incubated) towards the model dipeptide substrate S1 (Suc-LY-AMC) got enhanced by a 2.6-fold than its basal level activity without ADEP1 (Figure 3A). Similarly, on supplementation of bovine serum albumin (BSA) protein, enhancement in the peptidase activity of the ADEP1 bound rClpP1P2 (short-incubated) was detected; at the same time, neither the BSA nor its ClpP protomers degradation was noticed (Figure S2A and Figure S2B). In contrast, in the absence of the β-casein, a 7-fold reduction in the peptidase activity by the ADEP1 (40 μM) bound rClpP1P2 (short-incubated) was observed relative to its basal level activity without the ADEP1 (Figure 3A). On the other hand, the addition of ADEP1 (15 μM) to the pure ClpP isoforms failed to display any peptidase activity in the presence or absence of the β-casein (Figure 3B). Moreover, the addition of the β-casein substrate does not lead to a change in the measured peptidase activity in the absence of the ADEP1 (Figure 3B). The peptidase activity of the rClpP1P2 (long-incubated) bound to the ADEP1 demonstrated enhancement of the activity; at the same time, additional supplementation of the β-casein or BSA did not lead to any further gain in its activity (Figure S3A and Figure S3B). The supplementation of the β-casein to the ADEP1 activated rClpP1P2 (long-incubated) peptidase reaction abolished the self-cleavage of the ClpP protomers (Figure S3C). Additionally, we also examined if the ADEP1 bound rClpP1P2 (short-incubated) could perform the caseinolytic activity in a chaperone-independent process. Assessment of the ClpP caseinolytic reaction product on the denaturing polyacrylamide gel in the absence of the ADEP1 demonstrates that neither the pure rClpP isoforms nor their heterocomplex was able to degrade the β-casein or itself (Figure 4A, 4B and 4C, upper panels). Likewise, the presence of ADEP1 failed to demonstrate any caseinolytic or self-cleavage activity in the pure ClpP isoforms (Figure 4A and 4B, lower panels). However, the ADEP1 could trigger degradation of the β-casein (Figure 4C, lower panel) and the FITC-casein substrate (Figure 4D) by the rClpP1P2 (short-incubated) independent of the ATPase chaperone and without undergoing any self-cleavage.

**Figure 3.**
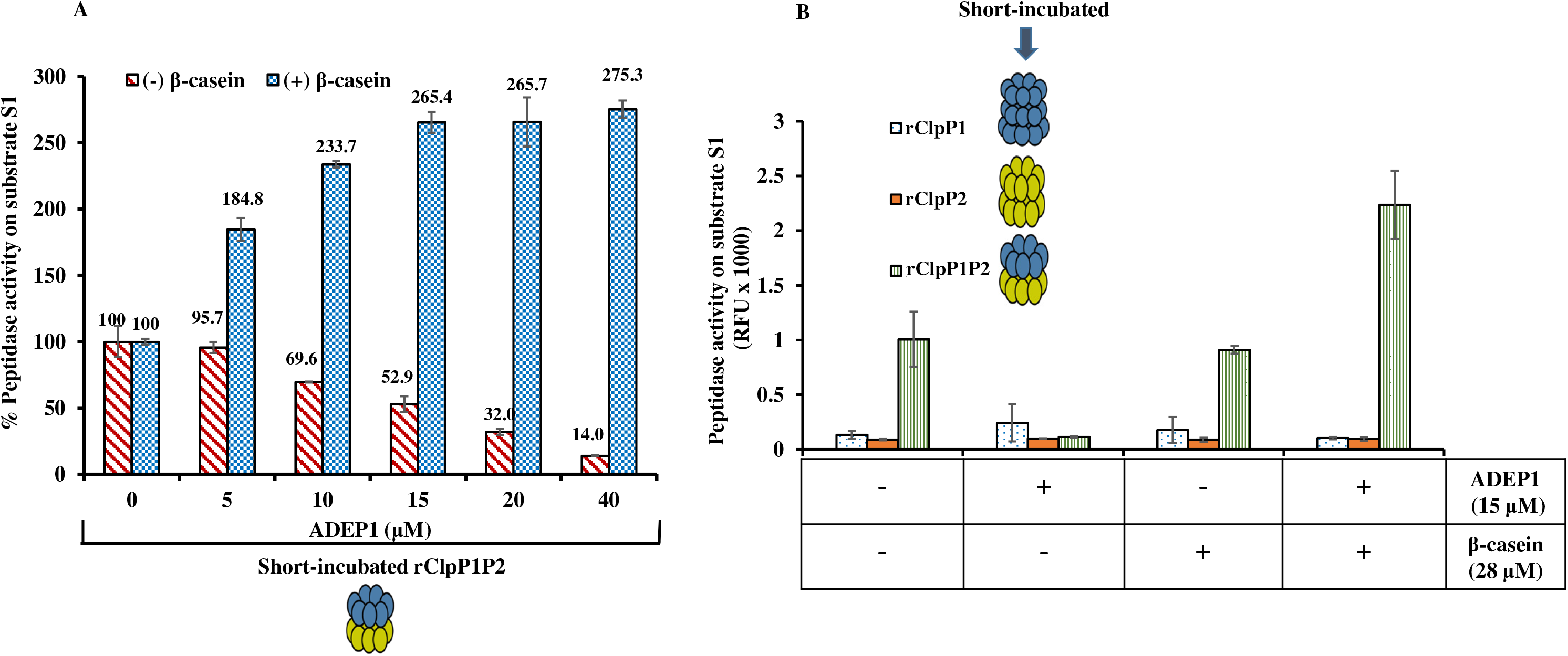
ADEP1 stimulated the peptidase activity of the rClpP1P2 (short-incubated) in the presence of the β-casein. **(A)** Comparison of peptidase activity of the rClpP1P2 stimulated by ADEP1 in the presence (+) or absence (−) of excess β-casein. Peptidase activity of the rClpP1P2 is represented as a percentage (%), wherein the end-point fluorescence was measured after 2 h of the enzymatic reaction. The measured end-point fluorescence value of the rClpP1P2 (containing no ADEP1) as control was considered as 100% for measuring the relative peptidase activity. For clarity, the schematics of the tetradecamer is shown. **(B)** Effect of the ADEP1 stimulated peptidase activity of the pure rClpP isoforms and its heterocomplex rClpP1P2 in the presence and absence of the β-casein. The error bars represent the standard errors of the mean (SEM) from the two-independent experiments performed.

**Figure 4.**
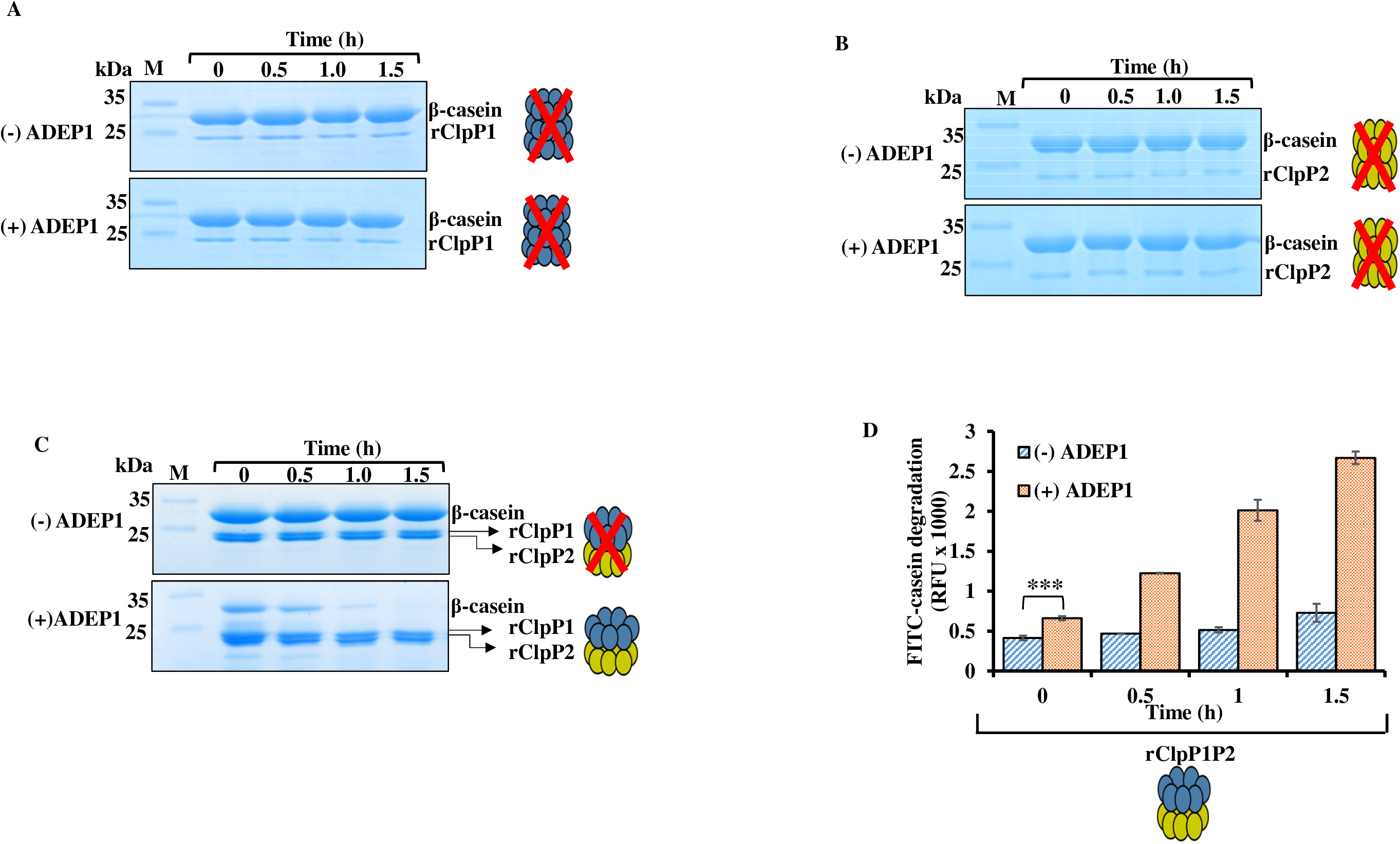
Effect of the ADEP1 on the protease activity of the pure rClpP isoforms and the rClpP1P2 heterocomplex of the *Leptospira.* **(A and B).** Denaturing gel electrophoresis showing the activity of the pure rClpP isoforms on the β-casein substrate in the presence (+) or absence (−) of the ADEP1. ADEP1 does not stimulate the pure rClpP isoforms (schematics in a red cross) for the degradation of the β-casein. **(C)** Denaturing gel electrophoresis showing the activity of the rClpP1P2 on β-casein in the (+) or (−) of ADEP1. The red cross over the schematics of the rClpP1P2 represents inactive heterocomplex. **(D)** Protease activity of rClpP1P2 after stimulation by the ADEP1 on the fluorogenic FITC-casein substrate. The error bars represent the standard errors of the mean (SEM) from the two-independent experiments performed (***p-value<0.001).

### ADEP1 exerts conformational influence over the whole tetradecameric rClpP1P2 of the *Leptospira*

Functional ClpP orthologs is a highly dynamic macromolecule (*12, 59*), and the proposed model states that the ClpP tetradecamer once stabilized remains in an equilibrium between an active (extended) and inactive (compressed) state (*26*). However, the ADEP1 bound ClpP orthologs smartly switches this equilibrium towards the extended state via conformational influence (*26*). So, we have strived to validate the compressed-to-extended state transition of the ADEP1 bound rClpP1P2 or the independent form of the tetradecamer by measuring the hydrodynamic diameter (D_h_) of the tetradecamer in a solution using the dynamic light scattering (DLS) technique as suggested before for the *Thermus thermophilus* ClpP (TtClpP) (*60*) and SaClpP (*37*). The measured diameter (D_h_) of the rClpP1P2 of the *Leptospira* in the presence (15 μM) and the absence of the ADEP1 were 16.76 and 13.60 nm, respectively (Figure 5A and 5B). The boost in the diameter of the ClpP machinery in the presence of the ADEP1 suggests an “open-gate” ClpP model that facilitates the entry of the unfolded substrates in the absence of the ATPase chaperone.

**Figure 5.**
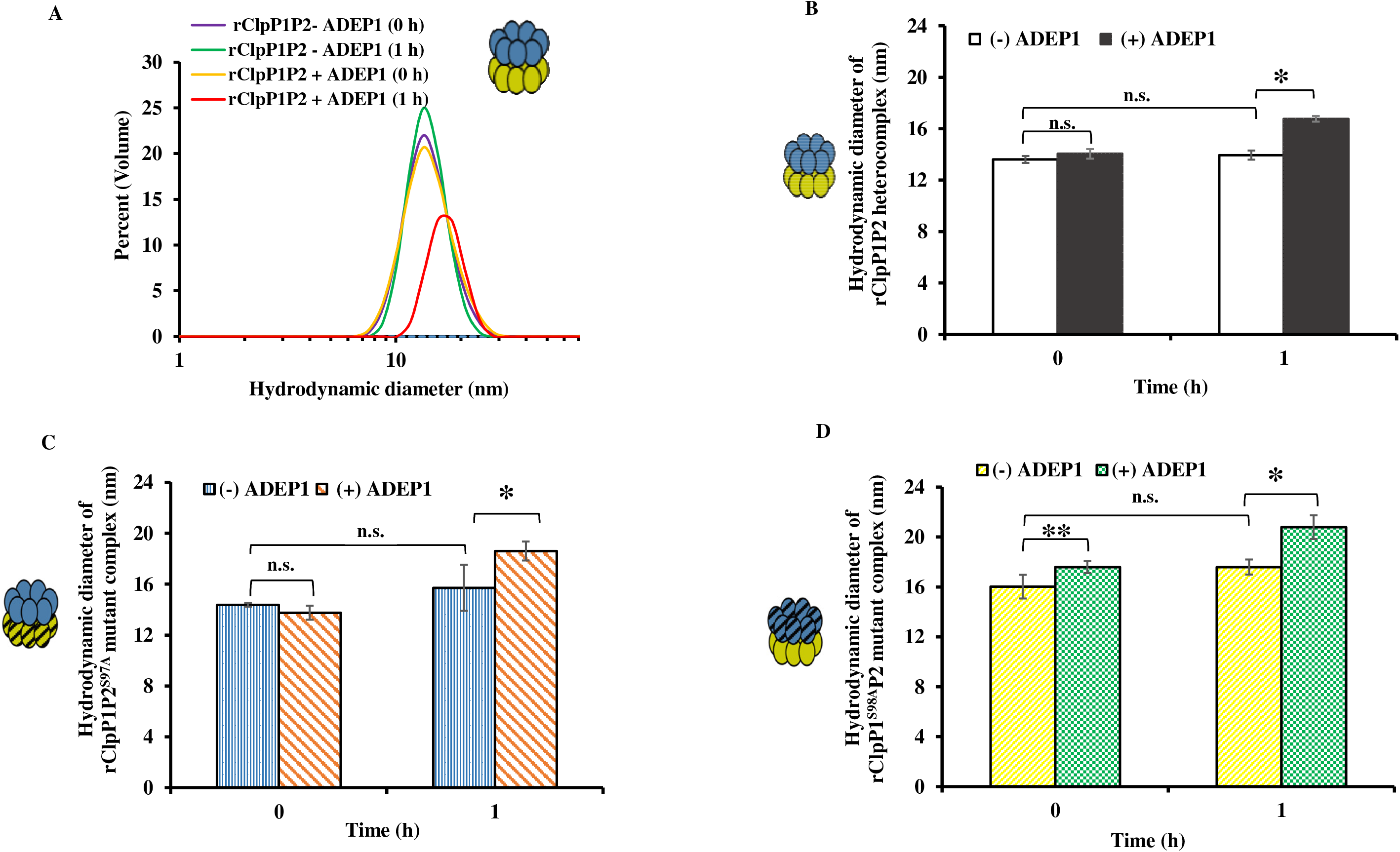
DLS analysis of the rClpP heterocomplex variants in the presence and absence of the ADEP1. **(A)** The representative DLS curves of the rClpP1P2 (0.5 mg mL^−1^) supplemented with (+) or without (−) the ADEP1 (15 μM). The relative volume (in %) versus particle size in nanometer (hydrodynamic diameter, D_h_) is plotted. The yellow and red curves represent the DLS curves of the rClpP1P2 in the presence (+) of the ADEP1 treatment at 0 h and 1 h, respectively. The violet and green curves represent the DLS curves of the rClpP1P2 in the absence (−) of the ADEP1 treatment at 0 h and 1 h, respectively. **(B)** Comparison of the average hydrodynamic diameter (D_h_) of the rClpP1P2 in the presence and absence of the ADEP1 at different time intervals (0 h and 1 h) using a bar graph. **(C)** Comparison of the average D_h_ of the mutant heterocomplex (rClpP1P2^S97A^) in the presence and absence of the ADEP1 at different time intervals (0 h and 1 h). **(D)** Comparison of the D_h_ of the mutant heterocomplex (rClpP1^S98A^P2) in the presence and absence of the ADEP1 at different time intervals (0 h and 1 h). The schematics with a diagonal line through them are the ClpP1^S98A^ (blue with black diagonal lines) and the ClpP2^S97A^ (yellow with black diagonal lines). The error bars represent the standard errors of the mean (SEM) from the two-independent experiments, where N = 15, the number of technical replicates for each of the protein complex in each of the experiments. Student’s t-test performed for statistical analysis to compare the measured D_h_ values (**p-value<0.005; *p-value<0.05; n.s. as not significant).

In our earlier study, we revealed that the mutant heterocomplex (rClpP1^S98A^P2 and rClpP1P2^S97A^) of the *Leptospira*, where the mutation of one of the catalytic triad serine (98/97) residue in the either of or both of the ClpP isoforms ushers to a loss of the peptidase activity. Moreover, the active site mutant variants of either isoform of the rClpP heterocomplex in association with its chaperone rClpX did not show any caseinolytic activity (*21*). The biochemical activity of the ClpP active site mutant variants in association with the ClpX implies simply an open-gate model of the ClpP is not self-sustaining for the gain of the caseinolytic activity.

It is established that the activation of the ClpP due to the binding of the ADEP1 relays various conformational changes in the whole ClpP machinery (*26*). This motivated us to address the question of whether the ADEP1 binding to the mutant heterocomplex (rClpP1^S98A^P2 and rClpP1P2^S97A^) steers to a perpetual increase in the D_h_ size as scaled for the rClpP1P2 heterocomplex of *Leptospira*? Forth DLS analysis, the measured structural diameter (D_h)_ of the rClpP1P2^S97A^ mutant heterocomplex in the presence (15 μM) and the absence of the ADEP1 were 18.60 and 14.38 nm, respectively (Figure 5C). On the other hand, the measured structural diameter of the rClpP1^S98A^P2 in the presence (15 μM) and absence of the ADEP1 were 20.79 and 16.03 nm, respectively (Figure 5D). The DLS of the serine mutant ClpP heterocomplex bound with the ADEP1 (15 μM) inferred a substantial expansion in the diameter (D_h_), where the difference was more striking in the rClpP1^S98A^P2 compared to the rClpP1P2 and the rClpP1P2^S97A^ variant (Table 1). The significant change in the structural diameter of the ClpP and its mutant variants in the presence of the ADEP1 indicates a conformational transformation in the whole ClpP machinery.

**Table 1.** Measured hydrodynamic diameter (D_h_) of the ClpP1P2 and its mutant heterocomplex variant in the presence (+) and absence (−) of the ADEP1.

| ClpP heterocomplex variants | Hydrodynamic diameter (nm) |  |
| --- | --- | --- |
|  | ADEP1 (-) | ADEP1 (+) |
| ClpP1P2 | 13.60±0.27 | 16.76±0.22 |
| ClpP1P2<sup>S97A</sup> | 14.38±0.14 | 18.60±0.75 |
| ClpP1<sup>S98A</sup>P2 | 16.03±0.95 | 20.79±0.96 |

### Serine^98^, a catalytic triad residue of the rClpP1 in the ClpP heterocomplex, is critical for the ADEP1 mediated activation

It has been hitherto unknown whether the two ClpP isoforms of the *Leptospira* are indistinctly susceptible to the ADEP1. The measured protease activity of the rClpP1P2 and its structural hydrodynamic diameter were more in the presence of the ADEP1 than in the absence of the ADEP1. Moreover, the ADEP1 bound rClpP1P2 is functionally a ClpX independent caseinolytic protease. This indirectly points towards the relaxed structural state of the rClpP1P2 in the presence of the ADEP1, which may arise due to the binding towards the apical surface of the ClpP hydrophobic pocket. Interestingly enough, the DLS of the serine mutant ClpP heterocomplex bound with the ADEP1 (15 μM), we witnessed the increase in the diameter (D_h_) to be more articulated in the rClpP1^S98A^P2 compared to the rClpP1P2 and the rClpP1P2^S97A^ variant. To explore the correlation of the increase in the diameter of the ClpP active site mutant variants of each isoform with its functional activity in the presence of the ADEP1, we aspired to gauge and compare its activity. Time-dependent casein (fluorescently labelled and unlabelled) degradation assay using the mutant ClpP heterocomplex (rClpP1P2^S97A^) bound to the ADEP1 demonstrated a gain in the protease activity (Figure 6A and 6B) in comparison to the mutant heterocomplex in the absence of the ADEP1. However, the ADEP1 bound rClpP1^S98A^P2 mutant heterocomplex failed to illustrate any gain in protease activity (Figure 6C and 6D), and so did the rClpP1^S98A^P2^S97A^ mutant heterocomplex (data not shown). Such differential biochemical and the biophysical properties of the reconstituted rClpP mutant heterocomplex within the active site variants of isoforms imply that the *Leptospira* ClpP displays an additional level of susceptibility towards the ADEP1, wherein the residue serine 98 of the ClpP1 is the preferred catalytic site of the ADEP1 interaction in comparison to the ClpP2 of the heterocomplex. These data further reflect that the ClpP1 active sites are more critical than the ClpP2’s in cleaving the model casein substrate and the active sites of the ClpP1 are a plausible location for the ADEP1 second interaction.

**Figure 6.**
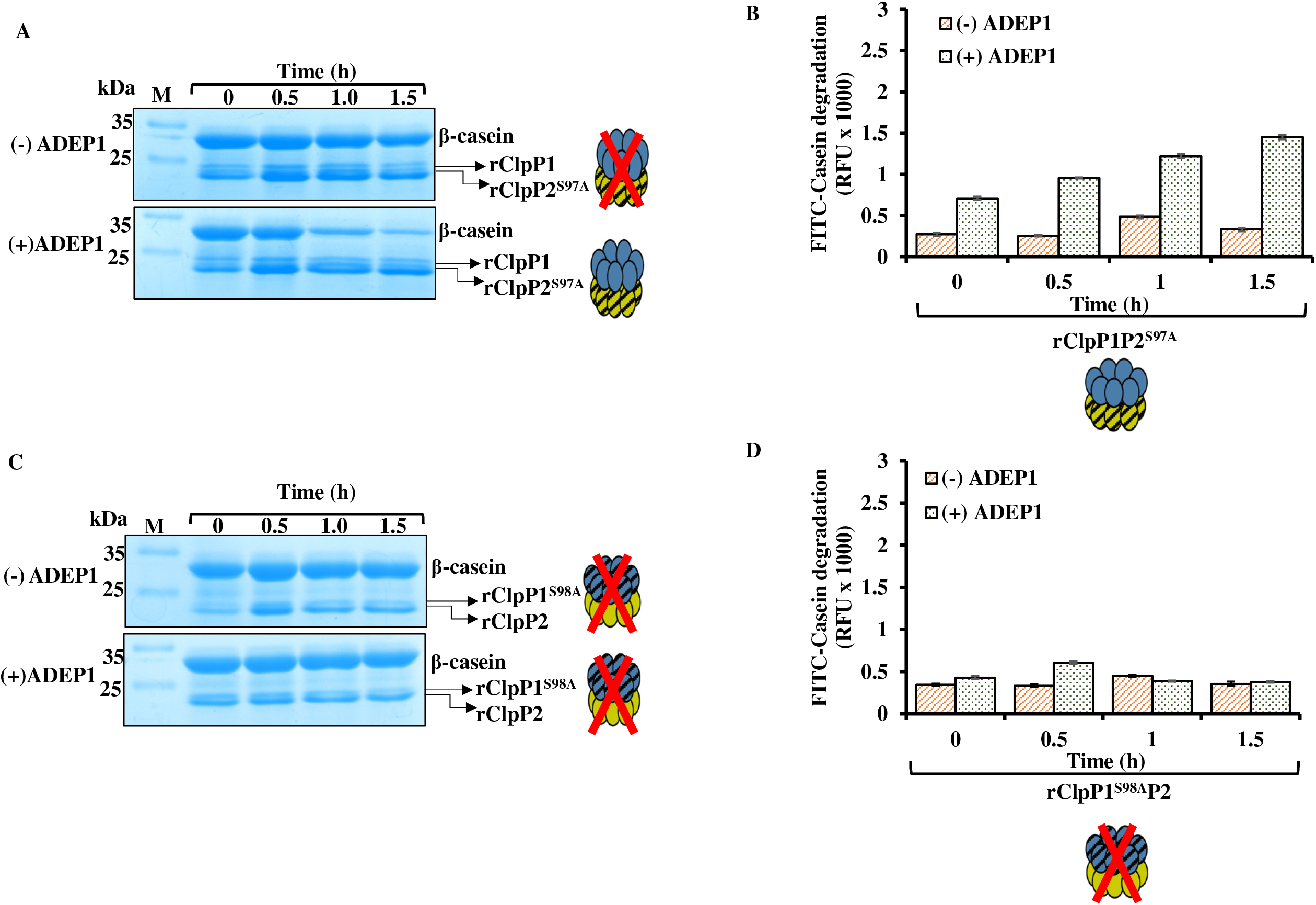
ADEP1 demonstrates predisposed interaction to the rClpP1 isoform of the *Leptospira* to configurationally switch on the ClpP machinery. **(A and B).**ADEP1 stimulates the protease activity of the mutant rClpP1P2^S97A^ heterocomplex on the β-casein and the fluorogenic FITC-casein substrates, respectively. With the increasing time of the ClpP protease reaction, the ADEP1 stimulates the mutant heterocomplex rClpP1P2^S97A^ to degrade β-casein (shown on the polyacrylamide gel stained with the Coomassie-blue) and the FITC-casein substrates (shown in fluorescence measurements). **(C and D)** ADEP1 fails to stimulate the protease activity of the mutant rClpP1^S98A^P2 heterocomplex on the β-casein and the fluorogenic FITC-casein substrates, respectively. The error bars represent the standard errors of the mean (SEM) from the two independent experiments performed.

### ADEP1 enhances the ClpXP1P2 complex activity of the *Leptospira*

We have illustrated previously that the leptospiral rClpXP1P2 complex is a caseinolytic protease in the presence of the ATP (*21*). Hence, we assessed the impact of the ADEP1 supplementation on the activity of the rClpXP1P2 complex of the *Leptospira*. The protease reaction illustrated a refinement in the degradation of FITC-casein by the leptospiral rClpXP1P2 complex in the presence of the increasing concentrations of the ADEP1 (5 - 20 μM), and afterward, the protease activity attained saturation (Figure 7A). Interestingly enough, without the aid of an ATPase chaperone (ClpX) and in the presence of the ADEP1 (5 - 20 μM), the rClpP1P2 showed more (4-fold) enhancement of the activity or can assert rampant stimulation than the rClpXP1P2 complex (Figure 7A). We also noticed a relatively lower protease enhancement of the rClpP1P2 than the rClpXP1P2 at the higher ADEP1 concentration (40 μM) implying the fallout of the ADEP1 on the rClpP1P2 differs at its higher concentration and the presence of the ClpX regulates the ClpP1P2 activity decently in the presence of the ADEP1 (Figure 7A). The justification for the straight enhancement of the rClpXP1P2 activity in the presence of the ADEP1 may be due to the simple increase in the axial diameter of the ClpP1P2 but it seems dubious as the ClpX and the ADEP1 compete for the same hydrophobic site. The second likelihood could be the ADEP1 interaction at the catalytic serine 98 residue of the ClpP1 (Figure 6A and Figure 6B), and the third possibility could be that ADEP1 interacts with the other unconventional site in the ClpP1P2 of the *Leptospira*. We formulated an independent protease assay to address the first two possibilities. We used the serine mutants rClpXP1P2 complex (rClpXP1^S98A^P2 and rClpXP1P2^S97A^) in the presence of an increasing concentration of the ADEP1 (5 - 40 μM) and compared with the protease activity of the rClpXP1P2 (Figure 7B). In consensus with our protease reaction embodied in figure 6, a reclaim of the protease activity could be detected exclusively in the mutant rClpXP1P2^S97A^ complex in the presence of the ADEP1, though it was lower than the rClpXP1P2 complex (Figure 7B). The protease reaction advocates that the regain in the protease activity of the mutant rClpXP1P2^S97A^ complex is predominantly due to the interaction of the ADEP1 at the catalytic serine 98 residue of the ClpP1 and not to that of the ClpP2 of the heterocomplex. A modest expansion in the axial pore diameter of the ClpP1P2 tetradecamer (by ClpX) could not activate the machinery. To further corroborate the analysis that the increase in the axial pore diameter of the machinery is not the solitary cause for an increase in the protease activity, an independent time chase protease experiment was executed. The time chase (0-1.5 h) protease assay using the mutant ClpP1P2^S97A^ in the presence of the ADEP1 (15 μM) was compared with the rClpXP1P2^S97A^ complex (Figure 7C). There was no statistically meaningful difference figured in the protease activity of both the ClpP1P2^S97A^ heterocomplex and the rClpXP1P2^S97A^ complex in the presence of the ADEP1 (Figure 7C). Collectively, the enhancement in the rClpP1P2 activity comes off to be because of the ADEP1 interaction to the active residue serine 98 of the ClpP1.

**Figure 7.**
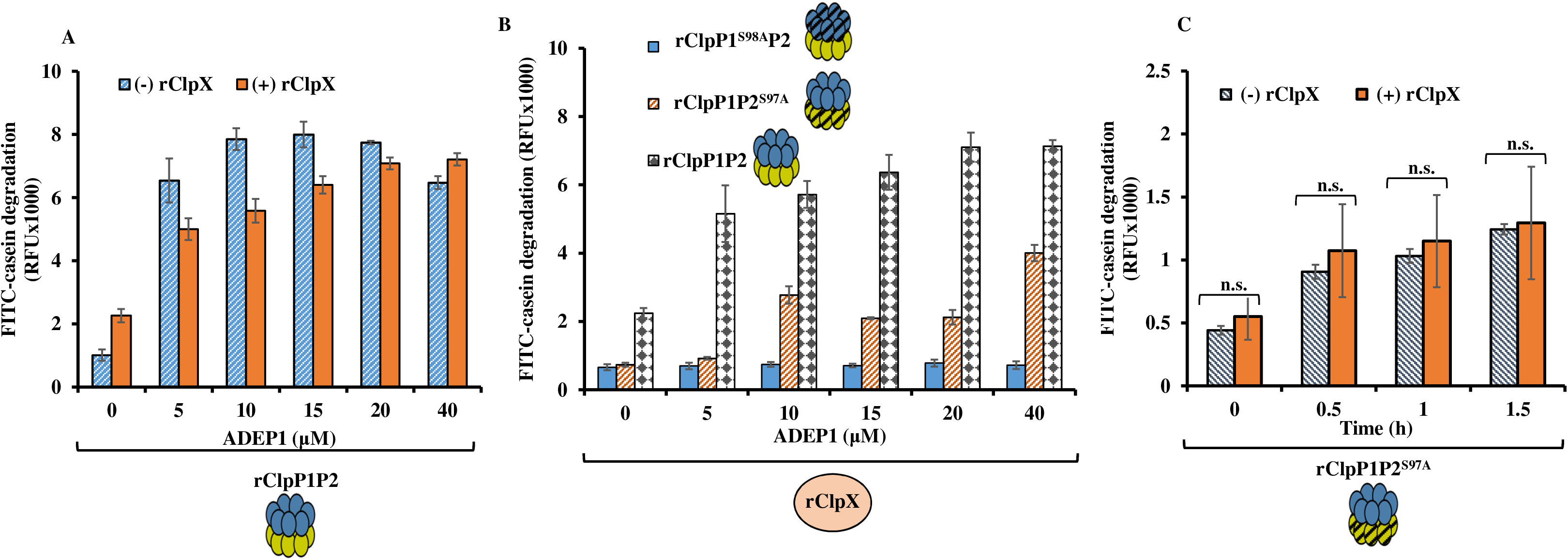
ADEP1 enhances the protease activity of the rClpXP1P2 and demonstrates biased catalytic activation of the mutant rClpXP1P2^S97A^ complex. **(A)** The substrate FITC-casein degradation (using fluorescence measurements) by the rClpP1P2 and the rClpXP1P2 complex in the presence of ADEP1. Increasing concentration of the ADEP1 (0 - 40 μM) enhanced the degradation of FITC-casein by the rClpXP1P2 complex. **(B)** Comparison of the protease activity of the rClpXP1P2 complex with that of the serine mutant’s complex (rClpXP1S98AP2 and rClpXP1P2^S97A^) in the presence of the ADEP1. The measured fluorescence in the absence of the ADEP1 for the rClp1P2 reflects the default background reading. **(C)** Comparison of a time chase (0-1.5 h) protease assay between the mutant ClpP1P2^S97A^ and the rClpXP1P2^S97A^ complex in the presence of the ADEP1. There was no significant difference statistically (p-value >0.05; n.s.) in the rate of protease activity between the mutant ClpP1P2^S97A^ and rClpXP1P2^S97A^ complex in the presence of an optimum concentration of the ADEP1 (15 μM). The error bars indicate the respective standard errors of the mean (SEM) from the two independent experiments performed.

### Model of the ClpP1P2 structure of *Leptospira* reflects the hydrophobic pocket lay in the ClpP2 subunits

In our earlier study, we showed that the various critical motifs of the ClpP isoforms in *Leptospira* crucial for conducting regulation are highly conserved in comparison to its known orthologs like the catalytic triad, Tyr activation trigger, Asp (Glu)/Arg oligomerization sensor domains, and the Gly-rich heptamer dimerization domain (*21*). In this study, the tertiary structure model of the ClpP1P2 tetradecamer of the *Leptospira* was developed using computational approaches. Although the ClpP1 and the ClpP2 subunits of the *M. tuberculosis* and the *L. interrogans* share a low sequence identity (~40%), their tertiary structures are very similar. A structural comparison of the ClpP1 and the ClpP2 subunits of the *Mycobacterium* and the *Leptospira* reveals that they are very analogous with an average root mean square deviation (rmsd) of ~0.6 Å. A representative model of the LepClpP1P2 complex along with the predicted potential ADEP1 binding hydrophobic pocket indicates a similarity between the *Mycobacterium* and the *Leptospira* (Figure 8A and Figure 8B). In theLepClpP1P2 heterotetradecamer model, the hydrophobic sites are exclusively present in the LepClpP2 heptamer apical region (Figure 8A and 8B). It is the hydrophobic binding sites of the LepClpP1P2 where the chaperone ClpX or the ADEPs may dock to constitute the operative protease. A comparison of the axial pore of the crystal structure of the ClpP1P2 complex of the *M. tuberculosis* with that of the modeled LepClpP1P2 structure shows that its diameter in former is throughout same from one to the other end while that in the later is conal in shape from the ClpP2 to ClpP1 end. Modeling of the LepClpP1P2 leads us to speculate that identical hydrophobic pockets as noticed in its orthologs are prevailing in the LepClpP2 required for the binding of its physiological chaperone (ATPase) or the antibiotic ADEP.

**Figure 8.**
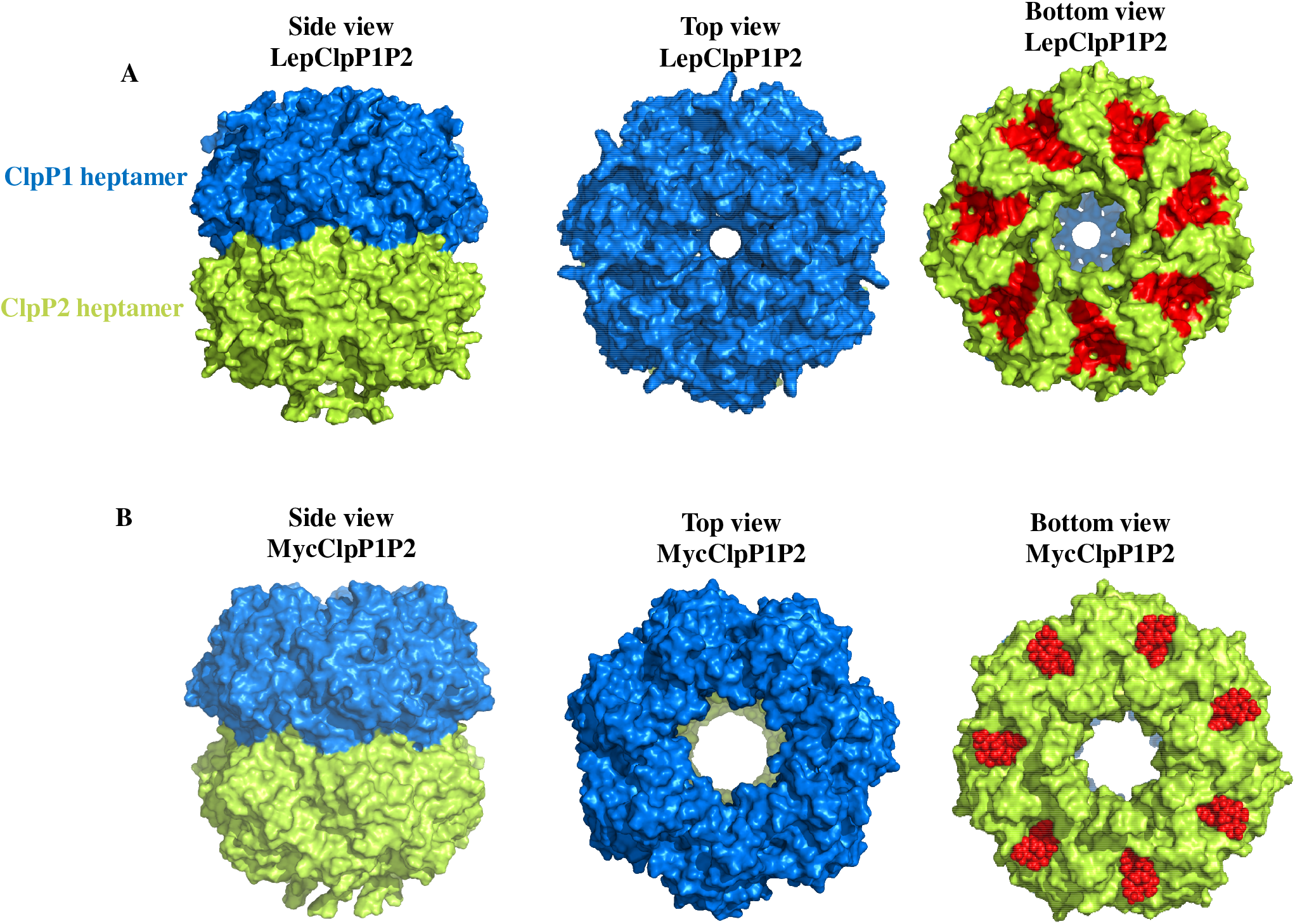
Modeling the tertiary structure of the ClpP1P2 of *Leptospira* and its comparison to the *Mycobacterium* ClpP. **(A and B).** The surface representation of the model ClpP1P2 heterocomplex from the *L. interrogans* and a crystal structure of the ClpP1P2 of the *M. tuberculosis*, respectively. The representative side, top, and bottom views of the *Leptospira* LepClpP1P2 are compared with the known crystal structure of the *Mycobacterium* MycClpP1P2, respectively. The ClpP1 and ClpP2 heptamer are represented in marine and limon color. The ADEP molecules bound to the MycClpP2 protein is represented as red spheres. A similar hydrophobic site at the LepClpP2 apical region to that of the ADEP binding of the MycClpP2 is highlighted in red, each spanning the two neighboring ClpP2 subunits at the apical surface.

## DISCUSSION

The Clp protease system as a target of antibiotic acyldepsipeptides (ADEPs) can be exploited for efficient bacterial killing either by the opening of the axial pore of the ClpP (*7, 10*) or by impeding the ClpP-ATPase chaperone interaction that disrupts regulated proteostasis (*12, 61, 62*). Nonetheless, the level of the susceptibility of the ClpP and its isoform towards the ADEP1 or its derivative are inconsistent as per the genus of the bacteria it belongs (*31, 55, 63*). In the current study, as proof of the concept, we chose to use natural antibiotic ADEP1 to investigate the molecular functioning of the ClpP protease of the spirochetes. The minimal inhibitory concentration (MIC) of the ADEPs for testing the *Mycobacterial* growth is in the range of 16 - 64 μg mL^−1^ (~23-90 μM) whereas in the *B. subtilis*, it is at a substantial lower range (nanomolar) (*12*). In this investigation, the *Leptospira* growth was interfered in the presence of 43 μg mL^−1^ADEP1 (60 μM) along with lengthening in its morphology. The eukaryotic cells are usually not affected by the ADEP1 up to the micromolar concentration range (*64*). Of note, in the *L. interrogans*, there is another closer ATPase dependent Clp protease (a threonine protease, ClpYQ, or HslUV), deletion of which ushered a failure of its survival in the hosts and the transmission of the leptospirosis (*65*). Consequently, the application of the ADEP1 for controlling the leptospirosis by targeting ClpP as an alternative to traditional drugs may be persuading. The alteration in the morphology of the *Leptospira* due to the antibiotic ADEPs has also been narrated in other bacteria like *Staphylococcus*, *Streptomyces*, and *B. subtilis* (*11, 35, 66*). Besides, in the *L. interrogans*, there are various other parameters (such as temperature, osmolality, host serum, stress) known to influence the gene expression and the spirochetes biology under *in vitro* conditions (*67*–*70*). Hence, understanding the *in vitro* impact of ADEP1 on spirochetes along with the elements that partly mimics ‘mammal-specific environments’ may be more insightful and persuasive.

To date, the investigation on the role and regulation of the Clp system in bacteria with more than one *clpP* gene by ADEPs is restricted to *M. tuberculosis* (*12, 26, 71*), *C. difficile* (*63*) and *Chlamydia* (*72*). The documented impact of ADEP on the multiple ClpP isoforms has been inconsistent under *in vitro* conditions. For instance, a synthetic ADEP could stimulate the peptidase activity specifically to only one pure ClpP isoform (ClpP1) of the *C. difficile* (*63*). On the other hand, in this study ADEP1 did not turn on any of the pure rClpP isoforms of the *Leptospira* even though the pure isoforms have the potential to oligomerize in the absence of the ADEP1 (*21*). The natural ADEP1 or its synthetic derivative has been established to activate the peptidase activity in both the single and multiple variants of the ClpP orthologs (*7, 12, 37*). Thus, in this study, the relative inhibition in the peptidase activity of the operative rClpP1P2 (short-incubated) in the presence of the ADEP1 was unanticipated. On the contrary, rClpP1P2 (long-incubated) despite the fact of encountering self-cleavage of ClpP subunits exhibited a relative gain in peptidase activity in the presence of the ADEP1 than the basal activity. With the numerous shreds of information obtained in this investigation and from the previous analysis (*21*), the relative decline in peptidase activity of ClpPs (short-incubated) in the presence of ADEP1 could be due to the self-cleavage of the ClpP protomers or the availability of a minor population of the stable and operative ClpPs or the cumulative effect of both factors. It is striking that on supplementation of the casein or BSA protein to the rClpP1P2 (short-incubated), the peptidase reaction in the presence of ADEP1 detected reversal in the inhibitions (2.6-3.4 fold activation) in relation to the basal activity without the ADEP1. The ClpP peptidase activity is directly proportional to the number of stable and operative self-assembled ClpP machinery. This is substantiated in a quite identical ClpP protease/peptidase experiment (Figure 4C, Figure S2B, and Figure S3C) wherein the supplementation of casein or BSA resulted in abolition in the self-cleavage of the ClpP protomers by the ADEP1 activated operative rClpP1P2 (short-incubated).

During the short-incubation period of the ClpP isoforms of the *Leptospira*, the dynamic equilibrium state between the operating heterotetradecamer ClpP machinery and its free ClpP protomers, there is a minor number of the operating stable ClpP peptidase machinery (*21*). It is established that the ADEP1 binds to the hydrophobic pocket on the outer edge of the apical surface of the ClpP, and once engaged at the interface between the adjacent monomers, it yields an incitation and the widening of the apical pore of the ClpP protease (*35*–*37*). In the *Leptospira* ClpP, the operative ClpP machinery (short-incubated) gets triggered by the ADEP1, and due to an increase in the diameter of the apical substrate channel, degradation of its free ClpP (more abundant) protomers may transpire. The stemming degradation of its ClpP subunits may veer the dynamic equilibrium further towards the free subunits as the short-incubated ClpPs are not very stable. Consequently, it may lead to a decline in the operative rClpP1P2 (short-incubated) peptidase machinery. Our assumption is in a consensus with an earlier investigation (*21*), where the binding affinity between the two rClpP isoforms of the *Leptospira* was moderate in range (K_d_,=2.02 ± 0.1 μM). Consistent with our observation in *Leptospira* ClpPs, numerous other time-dependent structure stabilization of the ClpP orthologs have been debated elsewhere (*12, 63*). The ClpP2 of the *Clostridium* assembled into a tetradecameric complex at ≥48 h of incubation (*63*) whereas, an incubation time of 4 h was a prerequisite to display catalytic activity in the ClpP1P2 of *Mycobacterium* (*12*). The self-cleavage of the ClpP subunits in the presence of the ADEP1 has also been documented in other bacteria like *B. subtilis*; regardless, there was a gain in the peptidase activity, and the cleavage was restricted to the 37 amino acids of its N-terminal (*7*).

Moreover, perhaps the influence of the ADEP1 is not equally binding in the case of an operative rClpP1P2 (long-incubated) wherein a surplus number of the stable and operative tetradecamer population is recorded (*21*). In the same study, the total peptidase activity of the rClpP1P2 (long-incubated) was higher than the rClpP1P2 (short-incubated) at a given time point (*21*). In concordance, in this analysis the total peptidase activity of the rClpP1P2 (long-incubated) was more enhanced with the ADEP1. The stimulation fallout of the ADEPs on the ClpP is also dangling upon other circumstances like the form of the ADEP used as an activator, catalytic acceleration at a ClpP serine residue and on the structural stability of the self-assembled ClpP tetradecamer (*36, 37, 63, 71*). In consensus to this, despite the ADEP1 mediated self-degradation of the ClpP protomers in the rClpP1P2 (long-incubated), there was a relative gain in the peptidase activity than the basal activity. The ADEP1 mediated rClpP1P2 (long-incubated) peptidase activity enhancement is in concordance to the earlier analyses on the ClpP orthologs, elsewhere (*7*). This investigation hence furnishes directives that there may be a second catalytic activation of the stable ClpP1P2 tetradecamer (long-incubated) by the ADEP1 that supersedes the influence of self-degradation. The catalytic activation of the ClpP by the ADEP1 is documented in *S. aureus* with the aid of chemical probes where the ADEP1 stimulates the ClpP activity through the cooperative binding (*37*). In the same study, it is suggested that the ADEP1 in addition to the opening of the ClpP axial pore by occupying the hydrophobic pocket, there is a ClpP conformational impediment into a more active form. On the same line, the DLS exploration of the ClpPs of the *Leptospira* in the presence of the ADEP1 ascertained us to accomplish conformational changes into a relaxed and active state. A site-directed mutagenesis analysis on the *Mycobacterium* ClpPs (ClpP1P2^S110^) possessing a mutation in the ClpP2 active site serine (Ser110), the addition of the ADEP analog resulted in the salvage of its protease activity. On the other hand, the *Mycobacterium* ClpPs (ClpP1^S98^P2) with a mutation in the ClpP1 active site (Ser98), the ADEP addition, could not support in retaining its activity (*31*). Likewise, in this study, it is apparent that the catalytic activation by the ADEP1 happens to be prejudiced towards the ClpP1 of the *Leptospira* as a mutation in the serine 98 residue stemmed in the complete abolition of the protease activity. At the same time, ADEP1 binding to the ClpP1P2^S97A^ in the *Leptospira* displayed retention of the activity. While it is to be pointed out that the ADEP1 mediated ClpP1 activation of catalytic serine seems to be equally important to the *Mycobacterium* and the *Leptospira* ClpPs, there is a conditional extra peptide agonist required for the ClpPs activation in the *Mycobacterium* (*31*). The catalytic activation or a gated-pore process activation of the ClpPs due to the ADEP has been noted even in multimeric compartmentalized proteases other than the Clp proteases (*73*–*75*). Lately, a proteasome inhibitor bortezomib has also been directed to bind to the ClpP active-sites serine of the *Thermus thermophiles*, emulating a peptide substrate and, evokes activity in the complex (*60*). Another accepted ClpP peptidase inhibitor β-lactones which can be used to bind specifically to the ClpP catalytic triad of the *Staphylococcus* to abolish the activity of the ClpP. Yet, the inhibitor errs to abolish the activity in the presence of the ADEP analog (ADEP7) (*37*).

Antibiotic ADEPs assist as an exploratory tool to reprogram the ClpP by a highly regulated peptidase to an autonomous and unregulated protease towards the unstructured proteins and the larger polypeptides, illustrated elsewhere (*35, 76*). It is suggested that a total of seven to fourteen ADEP molecule binds to the ClpP complex strictly where the ATPase chaperone engages with the functional ClpP tetradecamer (*54, 77, 78*). The number of ADEP1 molecules binding to the ClpP machinery relies on the availability of the hydrophobic pockets and whether the ClpP tetradecamer machinery is composed of two similar heptamers (ClpP1P1_7+7_) or with the two different homogenous heptamers (ClpP1P2_7+7_) stacked one above the other (*37, 54, 77*). The hydrophobic pocket thus signifies a hot spot for the ClpP modulation of the various organisms viz. *B. subtilis* (*76*), *Clostridium difficile* (*63*) and *E. coli* (*79, 80*). Interestingly, the *Mycobacterium* and the *Chlamydia* that encode two ClpP isoforms, the competent ADEP analogs bind to the seven hydrophobic pockets of the ClpP2 heptamer rather than the ClpP1 of the functional tetradecamer (*54, 71, 77*). The *Mycobacterium* ClpP1P2 binding site for the ADEP is correlated with the modeled ClpP1P2 of the *Leptospira* in this study. The computationally derived tertiary structure of the ClpP1P2 of the *Leptospira* infers that the ADEP1 can bind only to the hydrophobic pockets (7 in number) existing in the ClpP2 heptamer but not the ClpP1 heptamer like an operating ClpPs of the *Mycobacterium*. In the *Mycobacterium*, the biased ADEP binding towards the ClpP2 hydrophobic pocket of the operative ClpP heterocomplex also steers in simultaneous pore opening in the opposite ClpP1 ring implying structural interdependency within the ClpP machinery (*31, 34*). The axial diameter of the MycClpP1P2 in bound state with the ADEP and peptide agonist is indistinguishable from one end to the other while the modeled LepClpP1P2 structure shows the axial diameter of the complex from the ClpP2 to ClpP1 end conal in shape. Besides, the primary sequence alignment of the LepClpP2 with the MycClpP2 reveals to be shorter at its N-terminal end by eight residues whereas the ClpP1 is of proportionate size (*21*). Admittedly, it is too early to count on a model structure of the ClpP1P2 of the *Leptospira* unless the crystal structure of the ClpP1P2 bound to ADEP1 is developed.

In the genus, *Pseudomonas spp* that encode multiple ClpP isoforms, it is the ClpP1 that functionally interacts with the ClpX, whereas the ClpP2 exhibits no evidence of interaction to the ClpX (*81*) portraying the existence of a variant pattern of the Clp orthologs in nature. The measured gain in the activity of the ClpXP1P2 complex of the *Leptospira* in the presence of the ADEP1 in this analysis is in a consensus to the reported activity of the ClpAP complex of the *E. coli* in the presence of the ADEP1 (*35*). Regardless, in the *Mycobacterium* ClpP, the ADEP can competitively bind to the hydrophobic pocket of the ClpP2 heptamer and blocks of ClpX or ClpC1 engagement with the ClpP machinery leading to the abolition in the degradation of the natively folded protein GFP-ssrA or the unstructured casein substrate (*12, 31*). The ADEPs demolish many bacterial species by dysregulating the activity of the ClpP such that the multiple proteins are indiscriminately degraded (*35, 36, 76*), yet it annihilates the *Mycobacterium* by the inhibition of the essential ClpP-catalyzed proteolysis (*12, 31*). While drafting this paper, a surprising discovery caught our attention wherein a fragment of the ADEPs retained anti-Mycobacterial activity, yet stimulates rather than inhibits the ClpXP1P2-catalyzed degradation of the proteins (*71*). In the same study, they proposed the reasonable second interaction site at the ClpP1 catalytic triad for the ADEPs fragment. This explanation was in agreement with our finding, although the stimulation of the ClpXP1P2 in the *Leptospira* occurs by the full-length natural ADEPs and without any additional peptide agonists. Thus, the lethality of the *Leptospira* growth by the ADEP1 antibiotics can be assumed to be due to the enhancement of the native functions of the chaperone-dependent ClpP protease.

In this study, the relative higher caseinolytic activity of the ClpP1P2 of *Leptospira* in the presence of the ADEP1 than the ClpXP1P2 complex leads us to speculate that ClpX may not be dethroned entirely from the hydrophobic pocket by the ADEP1. This can be reasonably illustrated by the disparity in the mode of action of the ADEP1 and the ClpX. Despite being a competitor for the same apical hydrophobic pocket (*36*), ClpX is an ATPase dependent chaperone that would require time to unfold the substrate casein, whereas the ADEP1 works directly on the open-gate model. In one of the ClpXP proteolysis assay investigated elsewhere (*36*), approximately a 2-fold molar excess of the ADEP (calculated in relation to ClpP as a monomer) completely blocks the interaction of the ClpX with the ClpP of both *E. coli* and *B. subtilis*. In this investigation, in contrast to the *Mycobacterium* ClpXP, a consistent increase in the ClpXP activity of the *Leptospira* was noted in the presence of the increasing amount of ADEP1 implying towards advancement in indiscriminate degradation of the essential protein during the *Leptospira* growth.

A relative decline in the protease activity of the rClpP1P2 at a higher concentration of the ADEP1 (40 μM) than the rClpPXP1P2 may be explained in logic to the initiation of auto cleavage of the ClpP subunits. In contrast, in the presence of the ClpX, the ADEP1 mediated ClpP1P2 stimulation comes out to be more controlled. The precise explanation for the relative progress in the protease activity of the ClpXP1P2 in the presence of the ADEP1 is challenging to comprehend experimentally without any crystal structure. It is conceivable that the ADEP1 mediated gain in the protease activity may be due to the additional catalytic activation of the residue serine 98 of the ClpP1. To substantiate our understanding, we ascertained that the rClpXP1P2^S97A^ complex in the presence of the ADEP1 could be activated but not the rClpXP1^S98A^P2 complex. The expansion in hydrodynamic diameter was more pronounced for the rClpP1^S98A^P2 than the rClpP1P2^S97A^ complex in the presence of the ADEP1; nevertheless, mere gated-pore activation of the rClpP1^S98A^P2 complex did not transpire in any gain of the protease activity, thus reinforcing our postulate of the additional role of the ClpP1 active serine^98^ residue. Altogether, the different experimental studies on the ClpPs and its mutant in the presence of the ADEP1 establish a footing to investigate a more detailed portrait of the structure, function, and behavior of the ClpP system in the *L. interrogans*.

In our opinion, this is the first study about the impact of the ADEP1 on any pathogenic spirochete ClpP as an alternative to a regular antibiotic. Using the ADEP1 as a tool, this investigation provides an insight into the molecular function of the ClpP1P2 in a coalition with its ATPase chaperone ClpX of the *Leptospira*. Growth inhibition, biochemical assays, site-directed mutational analysis and *in silico* structure modeling of the ClpP1P2 gave a proof that the ClpP can be a suitable Achilles’ heel for the *Leptospira* by deregulating proteolysis inside the bacteria. The shreds of the evidence illustrated in this investigation verify that the antibiotic ADEP1 possesses the ability to control the ClpP system in a different approach than to the well-studied ClpP system in the *Mycobacterium*.

## METHODS

### Morphology changes and growth assays of the *Leptospira interrogans*

*L. interrogans* were grown *in vitro* at 29°C in 10 mL of the Ellinghausen-McCullough-Johnson (EMJH) medium supplemented with 5-fluorouracil till exponential phase. From the growing culture, 3×10^8^ cells were inoculated into the fresh EMJH media (1 mL) with or without the ADEP1 (10 μg mL^−1^ or 15 μM) dissolved in dimethyl sulfoxide (DMSO). The morphology of the cells was also investigated by assessing the length of the untreated and treated cells every 24 h till 120 h by the microscopy and imaging software (Zeiss). For generating the growth curve of the *L. interrogans* in the presence of different concentrations of ADEP1, an exponentially growing culture (100 μL containing 2×10^5^ cells) was added to a sterile non-binding white micro-test plate (96-well flat-bottom) in triplicate. Thereupon, to the cultures, ADEP1 was supplemented in the increasing concentrations (0, 20, 40, and 60 μM) and was incubated for 120 h at 29-30°C. The growth of the cells was monitored by counting the cell numbers on a hemocytometer counting chamber every 24 h till 120 h under the dark field microscopy (20× magnification). Each experiment was executed independently at least twice in triplicate.

### Field emission scanning electron microscopy (FESEM) of the *L. interrogans*

A 3 mL (6×10^7^ cells mL^−1^) of the exponentially grown culture of the *L. interrogans* in the EMJH medium with or without the addition of 43 μg mL^−1^ of ADEP1 (60 μM) was incubated till 48 h at 29°C. Post incubation, cultures were processed for the FESEM as illustrated before (*82*) with a few modifications. Briefly, the spirochetes were pelleted at 1500× g for 20 min, washed with phosphate buffer saline (pH 7.4), and fixed in glutaraldehyde (5% in 0.1 M phosphate buffer, pH 7.4) for 30 min at room temperature. Fixed samples were rinsed thrice with a phosphate buffer and dehydrated through a graded series of ethanol (35, 50, 75, 95, and 100% for 10 min each) followed by the final drying using hexamethyldisilazane (HMDS, 100%; Sigma) twice with a 10 min of incubation. Each time, cells were recovered by centrifugation. Over-night desiccated specimens (ADEP1 treated and untreated) were individually mounted on the aluminum stubs using double-sided carbon-coated tape, sputter-coated with the gold, and examined under the FESEM (Sigma-300, Zeiss, Germany) operated at 5 kV. The average length of a segment of the three complete spiral turns of ten spirochetes was measured to assess and correlate the partial length of treated and untreated *L. interrogans* in the representative of micrographs.

### Overexpression and purification of recombinant ClpP (rClpP) and rClpX of *Leptospira*

Caseinolytic protease (ClpP) isoforms and the chaperone ClpX of the *L. interrogans* serovar Copenhageni were cloned individually in the pET23a, overexpressed and purified from the *E. coli* BL21 (DE3) cells as illustrated before in our laboratory (*21*).

### Peptidase assays of *Leptospira* ClpP isoforms

The rClpP isoforms mixture (1.5-2 μg) were pre-incubated either for the 10 min at 37°C (short-incubation) or for the 24 h at 4°C (long-incubation) in a ClpP peptidase activity buffer (50 mM phosphate buffer pH 7.6, 100 mM KCl, 5% glycerol) to self-assemble into a functional heterocomplex. ADEP1 (BioAustralis, Cat No. BIA-A1570) was dissolved in DEPC-treated water with the 10% DMSO at a given working concentration (100 μM). ADEP1 was added at an increasing concentration (0-40 μM) into the flat bottom black polystyrene 96-well plates (Invitrogen) containing the rClpP heterocomplex and were incubated for 10 min at 37°C. Fluorogenic dipeptide substrate N-succinyl-Leu-Tyr-AMC (S1: Suc-LY-AMC; Sigma) was added (8 μL of 1 mM) to each of the wells to achieve a final substrate (S1) concentration (100 μM) in a given total reaction volume (80 μL). Assay plates were incubated for 2 h at 37°C, and the hydrolysis of the fluorogenic dipeptide was monitored via an i-TECAN Infinite M200 plate reader (excitation: 380 nm; emission: 460 nm). When using substrate β-casein in the peptidase activity assay of the short-incubated ClpP1P2, the same procedure was followed, as described above, with the supplementation of 28 μM of β-casein (Sigma) in the designated wells. Each experiment was performed at least twice in triplicates.

### Autoproteolysis assays of *Leptospira* ClpP isoforms

Pure rClpP isoforms (1.5-2 μg) or its mixture (short- or long-incubated rClpP1P2) into the ClpP peptidase activity buffer were incubated with a varying ADEP1 concentration (0-40 μM) in a given total reaction volume (20 μL) for 2 h at 37°C. Reactions were terminated by the addition of the sample buffer (SDS-PAGE loading buffer) and heating for 10 min at 95°C. The reaction products were resolved on the 12% SDS-PAGE and visualized by Coomassie staining.

### Protease assays of *Leptospira* ClpP isoforms

Pure rClpP isoforms or their mixture (2 μg) containing short- or long-incubated rClpP1P2 into a ClpP protease activity buffer (50 mM Tris-Cl pH 7.0, 50 mM KCl, 1 mM DTT, 8 mM MgCl_2_, 5% glycerol) was incubated with the ADEP1 (15 μM) for 10 min at 37°C. After the pre-incubation period, bovine β-casein (20 μM) was added to the reaction tube to a given final reaction volume (100 μL). From the total reaction volume, a given small volume (20 μL) of the reaction was terminated at the various intervals (0-1.5 h) after the addition of the sample buffer and heating for 10 min at 95°C. A control reaction of the pure rClpP isoforms or its mixture containing an equivalent amount of DMSO to the working solution of ADEP1, was included for comparison. The reaction products at each of the time points were resolved on 12% SDS-PAGE and visualized by Coomassie staining. A similar procedure was followed for the β-casein proteolysis assays wherein mutant isoforms of rClpP (rClpP1^S98A^ and rClpP2^S97A^) were used. In an alternative assay format, rClpP1P2, as well as its mutant isoforms, were evaluated for the protease activity in the presence of ADEP1 using fluorogenic substrate FITC-casein (Sigma). The rClpP1P2 or its mutant mixture (short-incubated) into the ClpP activity buffer was pre-incubated with the ADEP1 and was added to a 96-well black plate (Invitrogen). To each well, FITC-casein (10 μM) was added to a given final well volume (100 μL). The assay plates were then incubated in the dark for 2 h at 37°C, and the reactions were terminated with the trichloroacetic acid (0.6 N). Hydrolysis of the fluorogenic substrate was monitored via i-TECAN Infinite M200 plate reader (excitation: 492 nm; emission: 519 nm). Readings were obtained at every 0.5 h for 1.5 h. The protease activity of the rClpXP1P2 in the presence of the ADEP1 was measured using FITC-casein (10 μM) as the substrate in a given (100 μL) total reaction volume. In each of the reaction tube, the short-incubated rClpP1P2 (1 μg) were mixed with the rClpX (2 μg) and pre-incubated with a different ADEP1 concentration (0-40 μM) for 10 min at 37°C. After the pre-incubation period, 4 mM ATP was added to each of the reaction tubes to initiate the protease assay. The downstream of the assay was performed as described for the FITC-casein substrate degradation. To compare the protease activities of the ADEP-bound rClpXP1P2 complex and the mutant rClpXP1^S98A^P2 or rClpXP1P2^S97A^ complexes, similar FITC-casein degradation assay was carried out as described above. The activation of the rClpP1P2^S97A^ tetradecamer and the rClpXP1P2^S97A^ complex by the ADEP1 was measured by a time-chase degradation of FITC-casein. The mutant rClpP1P2^S97A^ (1 μg) and the rClpX (2 μg) were incubated shortly for 10 min at 37°C and mixed. ADEP1 (15 μM) was taken in a separate tube and further incubated with rClpP1P2^S97A^ or rClpXP1P2^S97A^ for another 10 min at 37°C. The reactions were initiated by the addition of FITC-casein (10 μM) and the ATP (4 mM) in a total reaction volume of 100 μL. From the total reaction volume, a given small volume (20 μL) of the reaction was terminated at various intervals (0-1.5 h) with the trichloroacetic acid (0.6 N). Hydrolysis of the fluorogenic substrate was monitored via i-TECAN Infinite M200 plate reader (excitation: 492 nm; emission: 519 nm). Each experiment was performed at least twice in triplicates.

### Dynamic light scattering

DLS experiments were performed on a Zetasizer Nano ZS (Malvern Instruments) at 25°C. Leptospiral rClpP1P2 or its mutant heterocomplex (ClpP1^S98A^P2 or ClpP1P2^S97A^) (0.5 mg mL^−1^) were incubated into a buffer (50 mM Tris-Cl pH 8.0, 100 mM NaCl and 10 % glycerol) for 48 h at 4°C for the self-assembly. The rClpP1P2 heterocomplex with or without ADEP1 (15 μM) was added to the polystyrene cuvettes to record the light scattering. The light scattering was recorded at 173° angle with a 633 nm He–Ne laser as the light source. The rClpP1P2 and the mutant heterocomplex were restored and further incubated for 1 h at 37°C, followed by the DLS of those samples. A total of 15 autocorrelation functions viz. technical replicates were recorded for each of the protein samples, and the hydrodynamic diameters were determined as described previously (*21*). Each investigation was performed in duplicate (2× for each the heterocomplex samples), and the hydrodynamic diameter was measured as the average of these replicates.

### Structure prediction of *Leptospira* ClpP

The tertiary structure models of the ClpP1 and ClpP2 from *L. interrogans* serovar Copenhageni were predicted using the web-based server Phyre2 (*83*). Subsequently, the predicted models were refined by the energy-minimization method using the program ModRefiner (*84*). *Leptospira* ClpP1, ClpP2, and ClpP1P2 oligomers were developed by superposing its monomers onto the known ClpP1P2 complex structures from the *Mycobacterium tuberculosis* (MycClpP1P2, PDB id: 4u0g). The stereo-chemical properties of all the refined models were validated using the webserver RAMPAGE (*85*). All the structural figures were prepared using the program PyMOL.

### Statistical analyses

All the results are expressed as means ± standard errors of the mean (SEM). Student’s paired t-test was used to determine the significance of differences between the means, and the p-values of <0.05 were regarded as statistically significant. At least two independent experiments were performed, each one in the duplicate or triplicate as mentioned in the materials and methods section and the figure legends.

## ACKNOWLEDGEMENTS

The authors gratefully acknowledge Dr Nitin Chaudhary, Department of Biosciences and Bioengineering, Indian Institute of Technology Guwahati (IIT Guwahati) for providing help in recording and analysing the DLS experiments. We acknowledge the Central Instruments Facility (CIF), IIT Guwahati for the FESEM.

## Author Contributions statement

MK conceived and supervised the study; MK and AD designed experiments and analyzed the data; SPK performed docking and modeling experiments; MSH performed the growth assays and the FESEM; MK, AD, and SPK wrote the manuscript.

