## Supplementary information for "ADEP1 activated ClpP1P2 macromolecule of *Leptospira*, an ideal Achilles’ heel to deregulate proteostasis and hamper the cell survival"

### Figure S1.

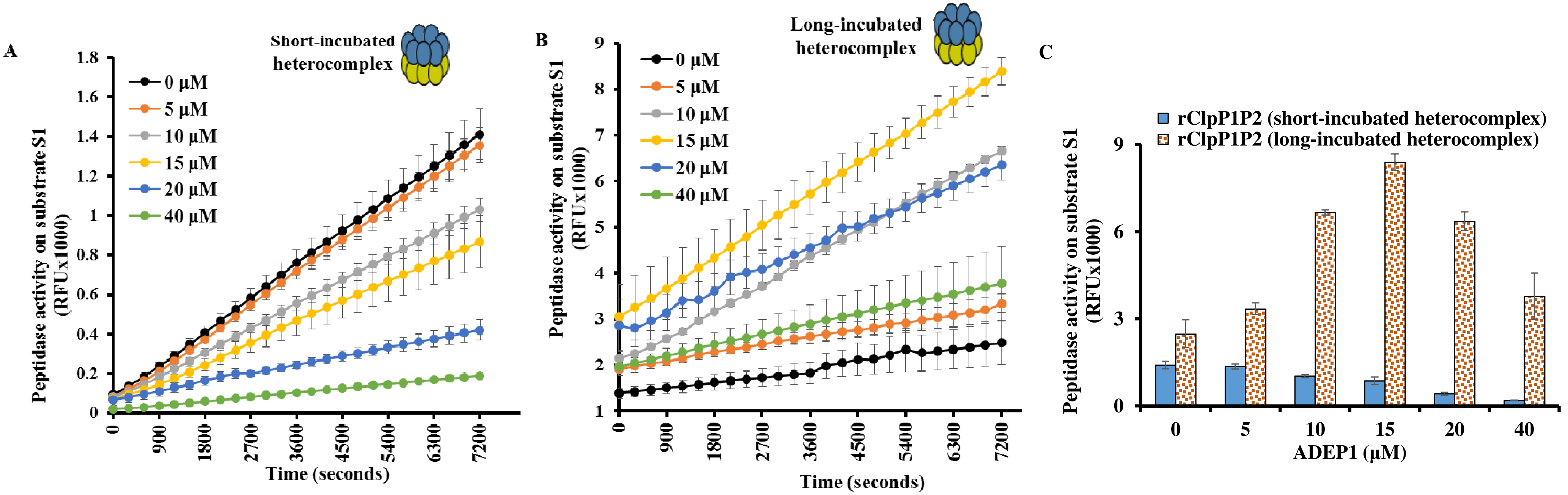

A
Short-incubated heterocomplex
0 µM
5 µM
10 µM
15 µM
20 µM
40 µM
Peptidase activity on substrate S1 (RFUx1000)
Time (seconds)
B
Long-incubated heterocomplex
0 µM
5 µM
10 µM
15 µM
20 µM
40 µM
Peptidase activity on substrate S1 (RFUx1000)
Time (seconds)
C
rClpP1P2 (short-incubated heterocomplex)
rClpP1P2 (long-incubated heterocomplex)
Peptidase activity on substrate S1 (RFUx1000)
ADEP1 (µM)

Figure S2

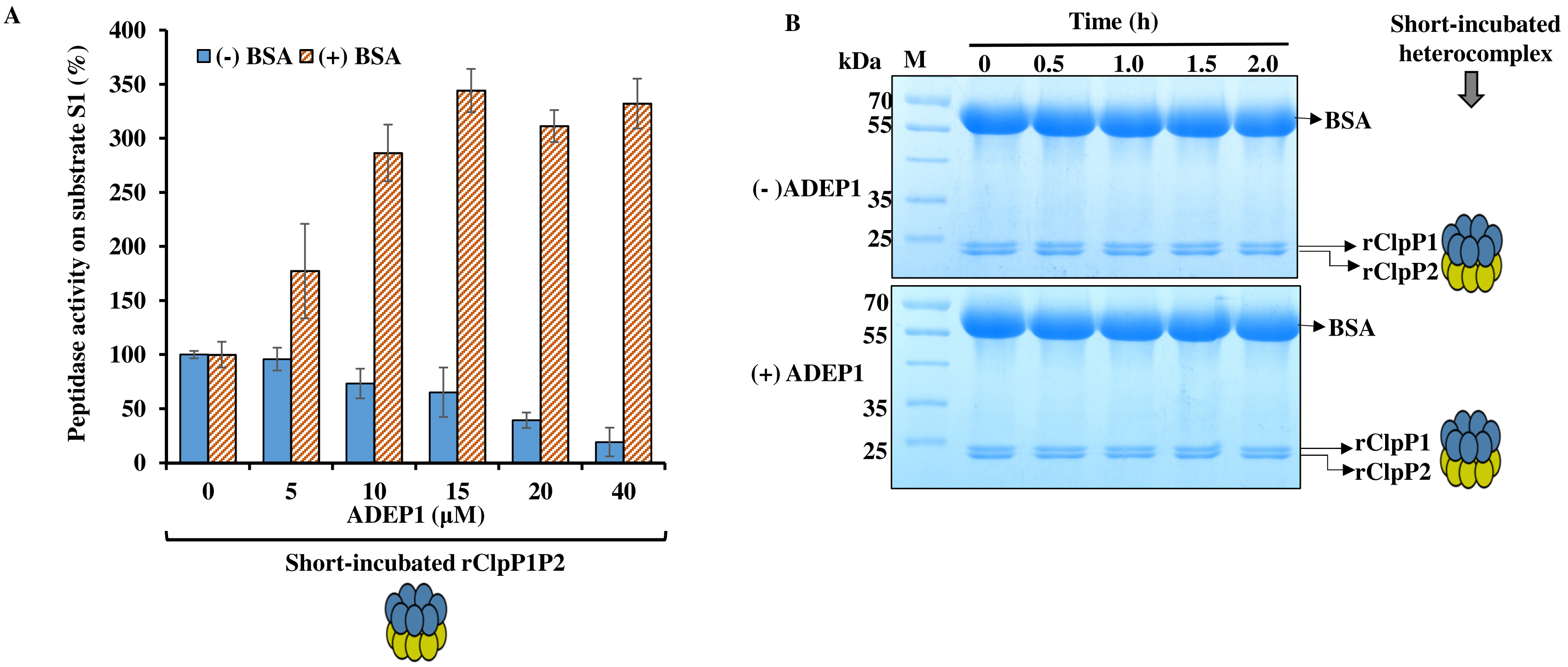
A
Peptidase activity on substrate S1 (%)
(-) BSA
(+) BSA
400
350
300
250
200
150
100
50
0
0
5
10
15
20
40
ADEP1 (μM)
Short-incubated rClpP1P2
B
Time (h)
kDa
M
0
0.5
1.0
1.5
2.0
Short-incubated heterocomplex
70
55
35
25
(- )ADEP1
BSA
rClpP1
rClpP2
(+) ADEP1
70
55
35
25
BSA
rClpP1
rClpP2

**Figure S3**

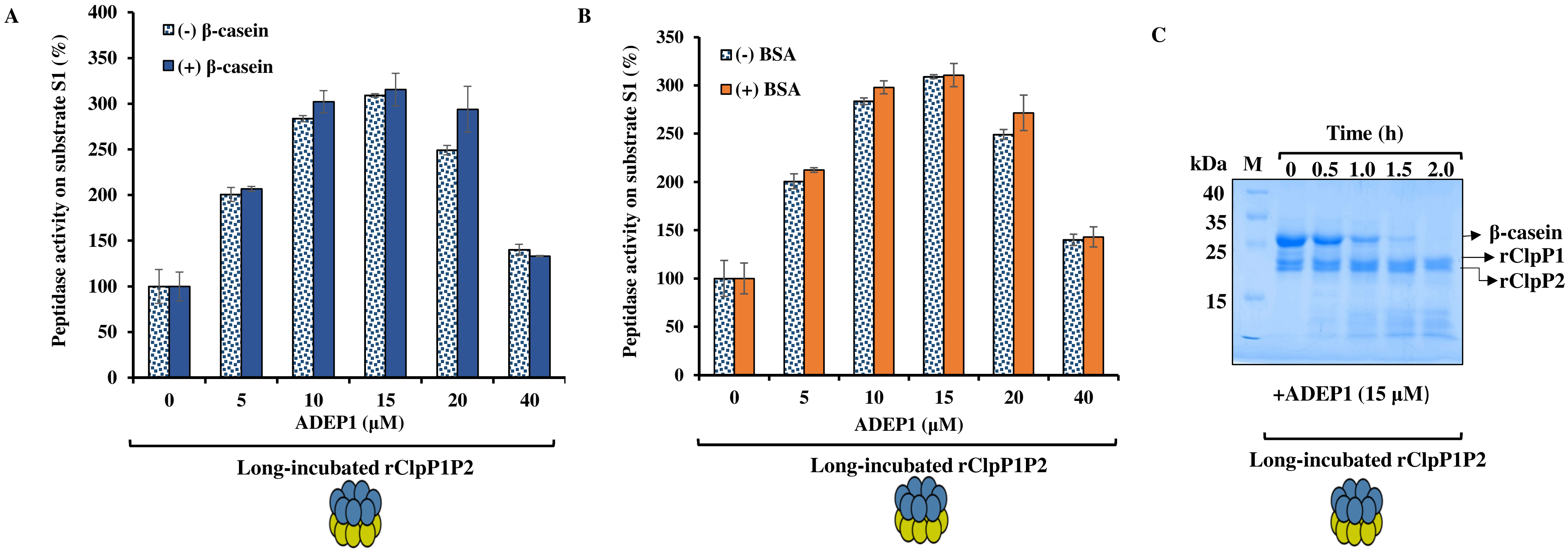

A
Peptidase activity on substrate S1 (%)
400
350
300
250
200
150
100
50
0
(-) β-casein
(+) β-casein
0
5
10
15
20
40
ADEP1 (μM)
Long-incubated rClpP1P2
B
Peptidase activity on substrate S1 (%)
350
300
250
200
150
100
50
0
(-) BSA
(+) BSA
0
5
10
15
20
40
ADEP1 (μM)
Long-incubated rClpP1P2
C
Time (h)
kDa
M
0
0.5
1.0
1.5
2.0
40
35
25
15
β-casein
rClpP1
rClpP2
+ADEP1 (15 μM)
Long-incubated rClpP1P2

### Supplementary Information

**ADEP1 activated ClpP1P2 macromolecule of *Leptospira*, an ideal Achilles' heel to deregulate proteostasis and hamper the cell survival**

Anusua Dhara, Md Saddam Hussain, Shankar Prasad Kanaujia, and Manish Kumar#

Department of Biosciences and Bioengineering, Indian Institute of Technology Guwahati,

Guwahati -781039, Assam, India

#corresponding author:

Manish Kumar

Department of Biosciences and Bioengineering, Indian Institute of Technology Guwahati, Guwahati-781039, Assam, India

Keywords: *Leptospira*, acyldepsipeptides (ADEP1s), Caseinolytic protease, ATPase, Peptidase, Casein

Running Title: ADEP1 activation of serine proteases of *Leptospira*

#### MATERIALS AND METHODS

##### Peptidase assays of *Leptospira* ClpP isoforms

The rClpP isoforms mixture (1.5-2 μg) were pre-incubated either for 10 min at 37°C (short-incubation) or for 24 h at 4°C (long-incubation) in ClpP peptidase activity buffer (50 mM phosphate buffer pH 7.6, 100 mM KCl, 5% glycerol) to self-assemble into functional heterocomplex. ADEP1 (BioAustralis, Cat No. BIA-A1570) was dissolved in DEPC-treated water with 10% DMSO at a given working concentration (100 μM). ADEP1was added at an increasing concentration (0-40 μM) into the flat bottom black polystyrene 96-well plates (Invitrogen) containing the rClpP heterocomplex and were incubated for 10 min at 37°C. Fluorogenic dipeptide substrate N-succinyl-Leu-Tyr-AMC (S1: Suc-LY-AMC; Sigma) was added (8 μL of 1 mM) to each of the wells to achieve a final substrate (S1) concentration (100 μM) in a given total reaction volume (80 μL). The hydrolysis of the fluorogenic dipeptide in the assay plates was monitored via an i-TECAN Infinite M200 plate reader (excitation: 380 nm; emission: 460 nm)at an interval of 5 min for 2h at 37°C reaction temperature. When using proteins β-casein or bovine serum albumin (BSA) in the peptidase activity assay of short- or long-incubated ClpP1P2, the same procedure was followed, as described above, with supplementation of 28 μM of β-casein (Sigma) or BSA (SRL) in the designated wells. Each experiment was performed at least twice in triplicates. The reaction products of short- and long-rClpP1P2 peptidase assay in the presence of 15 μM ADEP1 and supplemented with BSA/β-casein were withdrawn at various time-intervals (0- 2 h). The reactions were terminated by adding sample buffer and heating it for 10 min at 95°C. As ADEP1 is dissolved in the DMSO solution, a control reaction of rClpP1P2 containing an equivalent amount of dimethyl sulfoxide (DMSO) was set for comparison. The reaction products at each time point were resolved on 12% SDS-PAGE and visualized by Coomassie staining.

#### LEGENDS TO SUPPLEMENTARY FIGURES

**Figure S1. Absolute peptidase activity of the short-incubated rClpP1P2 in the presence of different ADEP1 concentration is lower than the long-incubated rClpP1P2. (A) and (B)** Peptidase activity of the short- and long-incubated rClpP1P2 on the dipeptide substrateS1 in the presence of a variable amount of ADEP1. Peptide degradation was measured fluorometrically as a relative fluorescent unit (RFU X 1000) at an interval of 5 min for 2 h. **(C)** Peptidase activity of the short- and long-incubated rClpP1P2 on the dipeptide substrate S1, where, end-point fluorescence was measured after 2 h of the enzymatic reaction. The error bars indicate the respective standard errors of the mean (SEM) from the two independent experiments performed.

**Figure S2. Peptidase activity of the ADEP1-bound rClpP1P2 (short-incubated) gets enhanced on the addition of BSA but the protease activity on BSA was not discernible. (A)** Comparison of peptidase activity of the rClpP1P2 stimulated by the ADEP1 in the presence (+) or absence (-) of the bovine serum albumin (BSA). Peptidase activity of the rClpP1P2 is represented as a percentage (%), wherein the end-point fluorescence was measured after 2 h of the enzymatic reaction. The measured end-point fluorescence value of the rClpP1P2 (containing no ADEP1) as control was considered as 100% for measuring the relative peptidase activity. ADEP1 increases the peptidase activity (~3.4-fold) of the rClpP1P2 in the (+) of BSA. For clarity, the cartoon representation of the ClpP tetradecamer used is presented. The error bars indicate the respective standard errors of the mean (SEM) from the two independent experiments performed. **(B)** Denaturing gel electrophoresis showing the activity of rClpP1P2 (short-incubated) on the BSA in the presence (+) or absence (-) of ADEP1. The resolved reaction products demonstrate the abolition of ClpP self-cleavage after the supplementation of BSA. BSA is not the substrate of chemoactivated ClpP1P2 but still prevents ClpP auto-degradation.

**Figure S3. The peptidase activity of the rClpP1P2 (long-incubated) bound to the ADEP1 demonstrated a gain of activity while supplementation of the β-casein or BSA did not lead to any additional stimulation in its activity. (A) and (B)** Comparison of the peptidase activity of the rClpP1P2 stimulated by the ADEP1 in the presence (+) or absence (-) of the β-casein/BSA. Peptidase activity of the rClpP1P2 is represented as a percentage (%), wherein the end-point fluorescence was measured after 2 h of the enzymatic reaction. The measured end-point fluorescence value of the rClpP1P2 (containing no ADEP1) as control was considered as 100% for measuring the relative peptidase activity.The error bars indicate the respective standard errors of the mean (SEM) from the two independent experiments performed. **(C)** Denaturing gel electrophoresis showing the reaction products (0-2hr) after stimulation of the peptidase activity of the rClpP1P2 (long-incubated) in the presence of 15 μM of the ADEP1 and supplementation of the β-casein. The rClpP1P2 (long-incubated) can degrade the unstructured model substrate β-casein with the abolition of the self-cleavage of its ClpP subunits.
